# Class I myosins direct circumferential F-actin flows to define cell chirality

**DOI:** 10.1101/2025.05.06.648335

**Authors:** Asuka Yamaguchi, Takeshi Sasamura, Takeshi Haraguchi, Kohei Yoshimura, Hiroki Taniguchi, Daisuke Kurisu, Yui Akano, Takamasa Higashi, Yasuhiko Sato, Florian L. Neugebauer, Mikiko Inaki, Yasuhiro Inoue, Masakazu Akiyama, Kohji Ito, Kenji Matsuno

**Affiliations:** Department of Biological Sciences, Graduate School of Science, The University of Osaka, 1-1 Machikaneyama-cho, Toyonaka, Osaka 560-0043, Japan; Department of Biology, Graduate School of Science, Chiba University, Chiba 263-8522, Japan; ZEISS Group, 2-10-9, Koji-machi, Chiyoda-ku, Tokyo 102-0083; School of Science, University of Hyogo, Ako-gun, Hyogo, 678-1297, Japan; Department of Micro Engineering, Kyoto University, Kyoto 615-8540, Japan; Department of Mathematics, Faculty of Science, Academic Assembly, University of Toyama, 3190 Gofuku, Toyama 930-8555, Japan; Membrane Protein Research Center (MPRC), Chiba University; Molecular Chirality Research Center, Chiba University; Plant Molecular Science Center, Chiba University

## Abstract

Eukaryotic cells possess intrinsic chirality in their structure, motility, and intracellular dynamics, which are designated cell chirality. Cell chirality participates in the left–right asymmetric morphogenesis and tissue integrity. However, the mechanisms of cell chirality formation remain elusive. In *Drosophila*, two evolutionarily conserved myosin I genes, *Myosin 1D* (*Myo1D*) and *Myosin 1C (Myo1C*), respectively, dictate the dextral and sinistral chirality of the cells and body. Here, we reported that Myo1D and Myo1C respectively directed the clockwise and counterclockwise circumferential flow of F-actin in *Drosophila* macrophages. Both induced the corresponding circular cytoplasm flows and depended on Myosin2 (Myo2). In a modified *in vitro* motility assay using near-physiological actin concentrations, Myo1D triggered the self-organization of the F-actin ring (chiral F-actin ring) that rotated clockwise; conversely, Myo1C induced the random flow of F-actin. The chiral F-actin ring implied that the F-actin bundle was parallelly and annularly polarized concerning its barbed pointed end. Considering that Myo1D and Myo1C are localized to the dorsal plasma membrane of macrophages, Myo1D and Myo1C might organize the parallelly polarized F-actin in macrophages. Our results suggest that Myo2 might drive the clockwise circumferential flow of F-actin along its parallel and annular polarity induced by Myo1D, which may be a molecular basis of cell and organ chirality.

## Main

Eukaryotic cells have intrinsic chirality in their shape, intracellular slewing motion, migration, and neurite extension, which are designated cell chirality^1–4^. Cell chirality is mostly described as the enantiomorphic states of these cellular features at the whole-cell level^5–7^. The chiral features of each cell also govern the placement of multiple cells under cultured conditions; consequently, cells become arranged in patterns with chirality^3,5,8,9^. Cell chirality is involved in the left–right (LR) asymmetric morphogenesis^1^ and the control of cell fates^10^, cancer metastasis^11^, tissue integrity, and permeability^12^. Studies have identified the factors required for the formation of cell chirality, such as actin polymerization regulators and actin crosslinkers, and consistent models have been proposed to show how they induce cell chirality^5,9^. In some cases, a given cell can show dextral (wild type) or sinistral (mirror image of wild type) cell chirality upon genetic modulations and changes in cellular conditions. For example, in *Drosophila*, *Myosin1D* (*Myo1D*) and *Myosin 1C* (*Myo1C*), both encoding evolutionarily conserved class I myosin, have dextral and sinistral activities, respectively, dictating the corresponding chirality of cells *in vivo*; consequently, they determine the dextral or sinistral LR asymmetry of various organs^1,2,13–15^. However, studies have yet to determine how a particular enantiomorphic state is selected and manifested at the whole-cell level. Additionally, Myo1D derives the chiral motion of a single F-actin molecule in an *in vitro* motility assay, suggesting that such molecular chirality consequently defines the dextral cell chirality^16^. Mouse myosin IC is known to induce chiral curved motion of single F-actin filaments similar to that observed with *Drosophila Myo1D* in *in vitro* motility assays^17^. A recent cryo-electron microscopy study revealed that this chiral motion arises from an oblique power stroke in which the lever arm of Myo1C moves at an angle relative to the actin filament axis during the pre-power stroke state, in contrast to conventional myosins whose lever arms swing parallel to the filament axis^18^. However, how the molecular chirality derived from these class I myosins is converted to cell chirality remains to be elucidated. Moreover, how Myo1C dictates sinistral cell chirality should be identified. To address these issues, we utilized the primary culture of *Drosophila* macrophages, which helped us conduct high-resolution time-lapse imaging to analyze the dynamics of F-actin^19^.

### *Myo1C* and *Myo1D* direct the circumferential flow of F-actin in cells

F-actin was visualized by the specific misexpression of *UAS-Lifeact-mGFP6* in macrophages isolated from the third-instar larvae and cultured on Concanavalin A-coated glasses^20^ (Fig. 1a and b). Their time-lapse movies were recorded at an interval of 15 s for 10 min (Fig. 1b). F-actin movement could not be attributed to the macrophage locomotion because they largely remained at the same position for 6 h (Supplementary Video1). As typically observed in cultured cells of various species^21^, Lifeact-mGFP6 exhibited a predominant retrograde F-actin flow (Fig. 1b and Supplementary Video 2). F-actin motility was quantified in macrophages via optical flow analysis^22,23^ and designated as the “direction vector of optical flow” (Fig. 1c). The direction vector of the optical flow of each pixel was split into two vectors: the unit direction vector to the direction of the cell centroid and the vector perpendicular to the unit direction vector to detect the chirality of F-actin motion (Fig. 1c). The latter represents the circumferential motion of F-actin around the cell centroid for 10 min in a clockwise (CW, minus value) or counterclockwise (CCW, plus value) direction, which was designated as the “chiral vector” (Fig. 1c). The magnitudes of the chiral vectors representing all pixels with positive and negative values in each cell were added, and the summation values were called the “chirality index” of F-actin flow. The chirality index obtained from wild-type macrophages (otherwise expressing *Lifeact-mGFP6*) was analyzed using a violin plot (Fig. 1d). Their chirality index showed a significant bias to CW (Fig. 1d and Table 1).

**Fig. 1.**
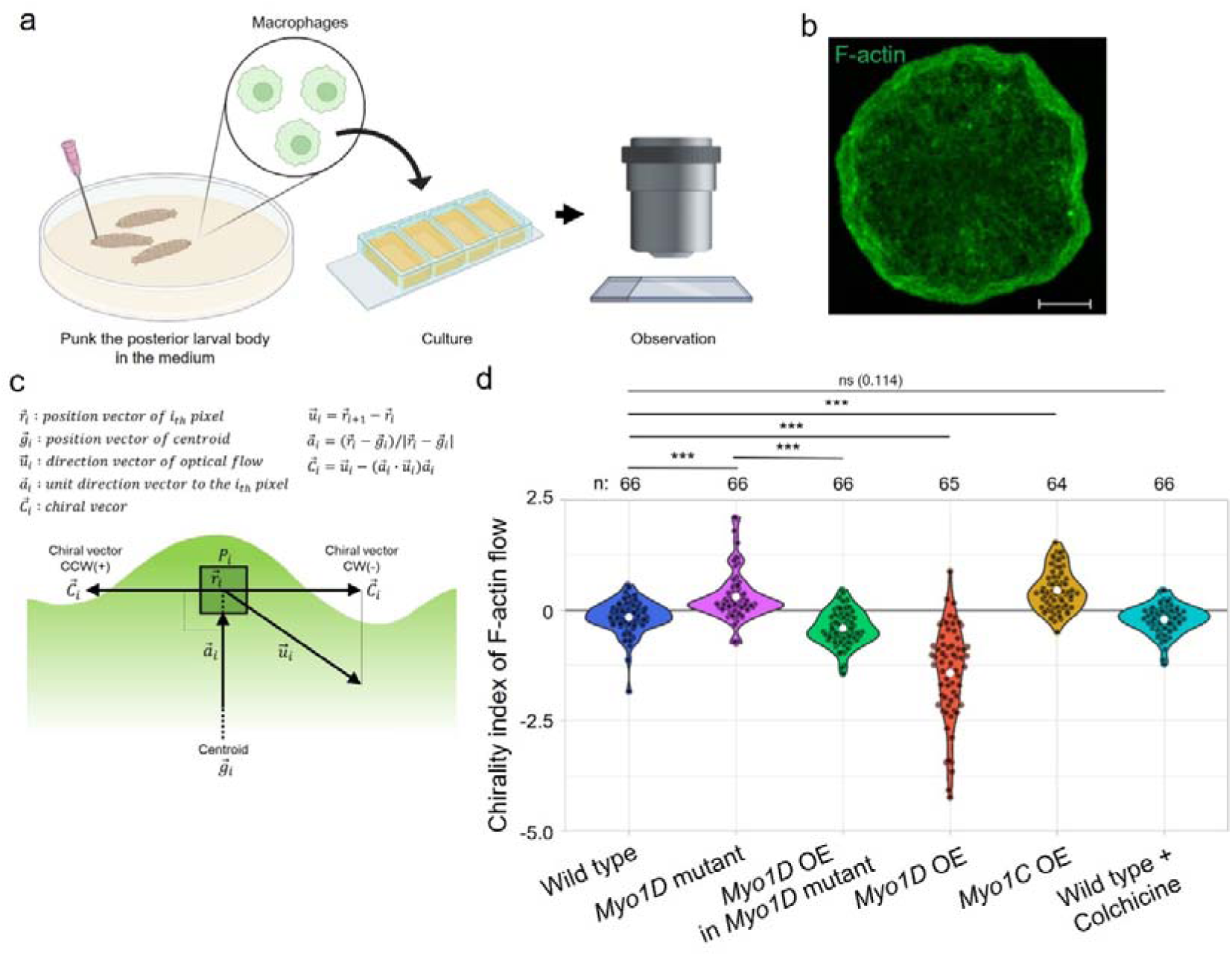
*Myo1D* and *Myo1C* induced CW and CCW rotations of F-actin, respectively, in *ex vivo* cultured macrophages. **a,** Procedure of live imaging to observe larval macrophages *ex vivo*. **b,** Time-lapse imaging of Lifeact-mGFP6 (green) visualizing the F-actin flow for 10 min. Scale bar is 5 μm. **c,** Green area schematically represents F-actin (b). F-actin flow was calculated by the optical flow estimation of each pixel, which is defined as a “direction vector of optical flow.” The direction vector of the optical flow was split into two vectors to reveal the chirality of F-actin flow: the position vector of the centroid (toward the cell centroid) and the “chiral vector” (perpendicular to the position vector of the centroid). The chirality of F-actin flow was defined by the chiral vector with CW (−) or CCW (+) rotational directionality. **d,** Violin plots showing the chirality index of F-actin flow (plus and minus values represent CCW and CW biases in the direction of F-actin rotation, respectively) in macrophages with genotypes and a treatment indicated at the bottom. White dots indicate the average values. The numbers of macrophages analyzed are indicated by “n” at the top of the graphs. The *P*-values were obtained by Steel–Dwass, and ns and *** represent *P* > 0.05 and *P* < 0.0001, respectively.

**Table 1.**
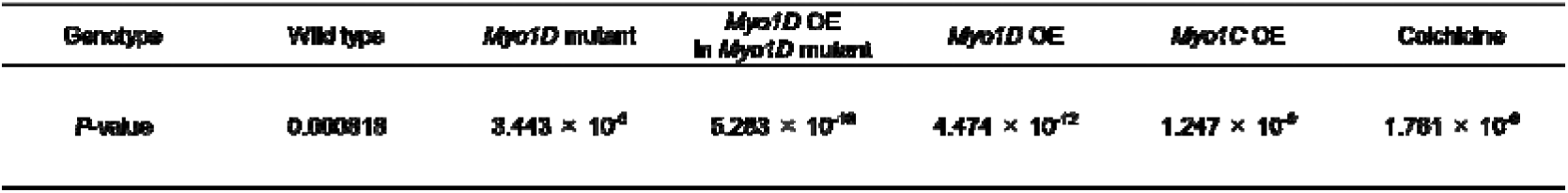
Statistical significance of CW or CCW biases in the chiral F-actin flow. The *P*-values of the statistical significance of CW or CCW biases in the chiral vector of the F-actin flow in the macrophages of the indicated genotypes and treatment were calculated via a one-sample one-sided Wilcoxon signed-rank test.

The activities of *Myo1D* and *Myo1C* were analyzed using the same procedures to direct the chirality of F-actin flow. Single-cell RNA sequencing analysis previously revealed that *Myo1C* and *Myo1D* in wild-type macrophages are endogenously expressed at low and modest levels, respectively^24^. In macrophages that are homozygous for a null mutant of *Myo1D*, their chirality index showed a significant CCW bias (sinistral), which was the mirror image of the wild-type counterpart, recapitulating its cell- and organ-chirality phenotypes *in vivo*^1,2^ (Fig. 1d and Table 1). The specific overexpression of wild-type *Myo1D* in *Myo1D* mutant macrophages effectively restored such CCW-bias to CW-bias with statistical significance (Fig. 1d and Table 1). Therefore, *Myo1D* mutation is responsible for the mirror reversal of the F-actin flow’s direction. Furthermore, the *Myo1D* overexpression in wild-type macrophages significantly enhanced the CW-bias because almost all macrophages showed a CW rotation (Fig. 1d and Table 1). Therefore, collectively, *Myo1D* exhibits the dextral activity to dictate the slewing direction of F-actin flow, which is consistent with the dextral roles of *Myo1D* in other cell types *in vivo*^13,16^. Although loss-of-function *Myo1C* alone subtly affects the LR asymmetry in *Drosophila*, its overexpression in wild-type flies efficiently reverses LR asymmetry in various organs^14,16^. In the present study, *Myo1C* was overexpressed in wild-type macrophages; their chirality index was significantly inversed from CW to CCW, demonstrating its sinistral activity in chiral F-actin flow (Fig. 1d and Table 1). Therefore, the enantiomorphic states of chiral F-actin flow at the whole-cell level are determined by *Myo1D* and *Myo1C*.

### Chiral F-actin flow is coupled with cytoplasmic dynamics with respective chirality

Considering that F-actin flow generally controls the dynamics of the cytoplasm, we speculated that chiral F-actin flow may rotate the whole cytoplasm, which may account for the change in the chiral cell shape observed *in vivo*^1,25^. In macrophages, the centrosome was found to be a clear landmark structure in the cytoplasm, which could be visualized using *Centrosomin (Cnn)-GFP*^26^ and tracked via time-lapse analysis with low phototoxicity (Fig. 2a, Supplementary Video 3). The circumferential motion of a centrosome around the center of the nucleus was represented by negative (CW) and positive (CCW) angle *θ* (degree) every 30 min for 6 h (Fig. 2b and c). The rotation angles *θ* of centrosomes at the time point after 6 h (*θ_fina1_*) were analyzed using a violin plot in wild-type macrophages (Fig. 2 d). Their centrosomes tended to rotate CW rather than CCW with statistical significance (Fig. 2d, Table 2, and Extended Data Fig. 1b). The correlation coefficient between the chirality index of F-actin (x-axis) and the centrosome rotation (angle *θ_fina1_*) (y-axis) was calculated to test a causal relationship between the chirality of F-actin flow and centrosome rotation, which were simultaneously observed in wild-type macrophages for 20 min (Fig. 2e). Their correlation coefficient (r=0.383) indicated that these two events could be positively correlated (Fig. 2e). Since the centrosome is a part of a microtubule network, we excluded the possibility that the microtubule network drives the chiral F-actin flow. We added colchicine to inhibit microtubule polymerization and examined its effect on the chirality index of F-actin flow in wild-type macrophages. Under the conditions in which microtubule formation was largely diminished (Extended Data Fig. 1a), our violin plot revealed that the significant CW-bias of the chiral F-actin flow was maintained, demonstrating that microtubules are inessential for the chiral F-actin flow (Fig. 1c, Table 1).

**Fig. 2.**
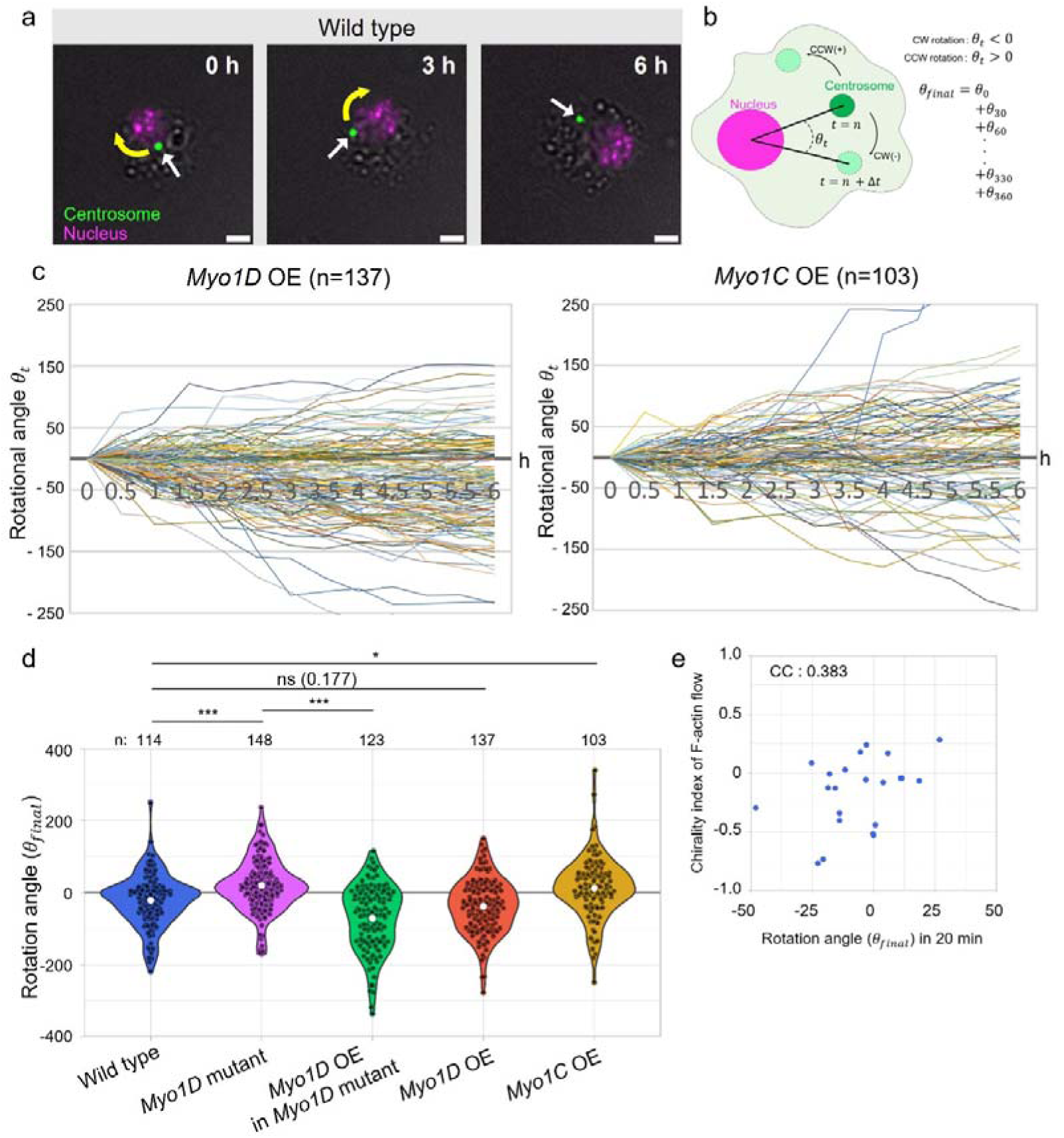
*Myo1D* and *Myo1C* induced the CW and CCW rotations of the centrosome around the nucleus, respectively, in macrophages. **a,** Snapshot of the centrosome (green, indicated by white arrows) and nucleus (magenta) labeled by Cnn-GFP and Redstinger (magenta), respectively, in wild-type macrophages at the time points indicated at the top right. Scale bars represent 5 μm. **b,** Schematic of the calculation of the rotation angle (*θ_t_*) of the centrosome movement from t=n (deep green circles) to t=n+Δt (light green circles with CW [−] or CCW [+] rotation) around the nucleus (magenta circle). “t” represents any time point from 0 min to 360 min. *θ_fina1_* represents the integrated rotation angle of the centrosome for 360 min. **c,** Rotation angles (*θ_t_*) of the centrosome from t=0 min at every 30 min (0.5 h) for six h (plus and minus values represent the CCW and CW direction of the rotation, respectively) in macrophages overexpressing *Myo1D* (*Myo1D* OE) or *Myo1C* (*Myo1D* OE). Polygonal lines shown in different colors represent the results of each macrophage. The numbers of the examined macrophages are shown in parentheses. d, Violin plots showing the rotation angle (*θ_fina1_*) of the centrosome (plus and minus values represent CCW and CW biases in the direction of the rotation, respectively) in macrophages with genotypes indicated at the bottom. White dots indicate the average values. The numbers of the analyzed macrophages are indicated at the top of the graphs. The *P*-values were obtained by Steel–Dwass, and ns, *, and *** represent *P* > 0.05, *P* < 0.05, and *P* < 0.0001, respectively. **e,** Correlation analysis between the chirality index of actin flow and rotation angle (*θ_fina1_*) for 20 min in wild-type macrophages. CC: Correlation coefficient assessed by Pearson correlation analysis.

**Table 2.** Statistical significance of CW or CCW biases in the rotational angle of the centrosome. The *P*-values of the statistical significance of CW or CCW biases in the rotational angle of the centrosome for 360 min ( ) in macrophages with the indicated genotypes were calculated via a one-sample one-sided Wilcoxon signed-rank test.

| **Genotype** | **Wild type** | ***Myo1D* mutant** | ***Myo1D* OE**<br>**in *Myo1D* mutant** | ***Myo1D* OE** | ***Myo1C* OE** |
| --- | --- | --- | --- | --- | --- |
| ***P*-value** | **0.00101** | **0.000488** | **$9.053 \times 10^{-13}$** | **$4.003 \times 10^{-8}$** | **0.0372** |

We examined the effect of *Myo1D* and *Myo1C* on the chirality of centrosome rotation. In the line graphs showing the rotation angles (*θ*) of each centrosome every 30 min for 6 h in *Myo1D*- and *Myo1C*-overexpressing macrophages, the densities of lines were more crowded in the CW and CCW areas, respectively (Fig. 2c). The macrophages of the *Myo1D* mutant without or with overexpressed *Myo1D* and macrophages overexpressing *Myo1D* or *Myo1C* exhibited the CW/CCW chirality of centrosome rotation, which coincided with the effects on their chirality index of F-actin flow in the respective genetic conditions of macrophages, as revealed by violin plots (Fig. 1d, Fig. 2d, Extended Data Fig. 1b, Table 2). Thus, *Myo1D* and *Myo1C* directed the CW and CCW chirality of centrosome rotation, respectively. Therefore, chiral F-actin flow defines the CW or CCW bias of intracellular dynamics.

### *Myo1D* and *Myo1C* expand the areas where F-actin shows CW and CCW chiral F-actin flow, respectively

To understand how the CW or CCW bias of chiral F-actin flow is induced in each cell, we investigated the intracellular distribution of the area where F-actin demonstrated the CW or CCW bias. In this analysis, we showed the value of F-actin chiral vectors in each pixel with the classification of CW (yellow) and CCW (magenta) for 10 min (Fig. 3a and Extended Data Fig. 2). We found that the distribution of F-actin chiral vectors with CW or CCW directionalities appeared in a mosaic pattern, which was composed of CW and CCW domains, referred to as “chiral F-actin domains” in this study (Extended Data Fig. 2). In all genotypes examined, the chiral F-actin domains appeared as scattered patterns of CW and CCW domains but not the monopolization of either of the domains (Extended Data Fig. 2).

**Fig. 3.**
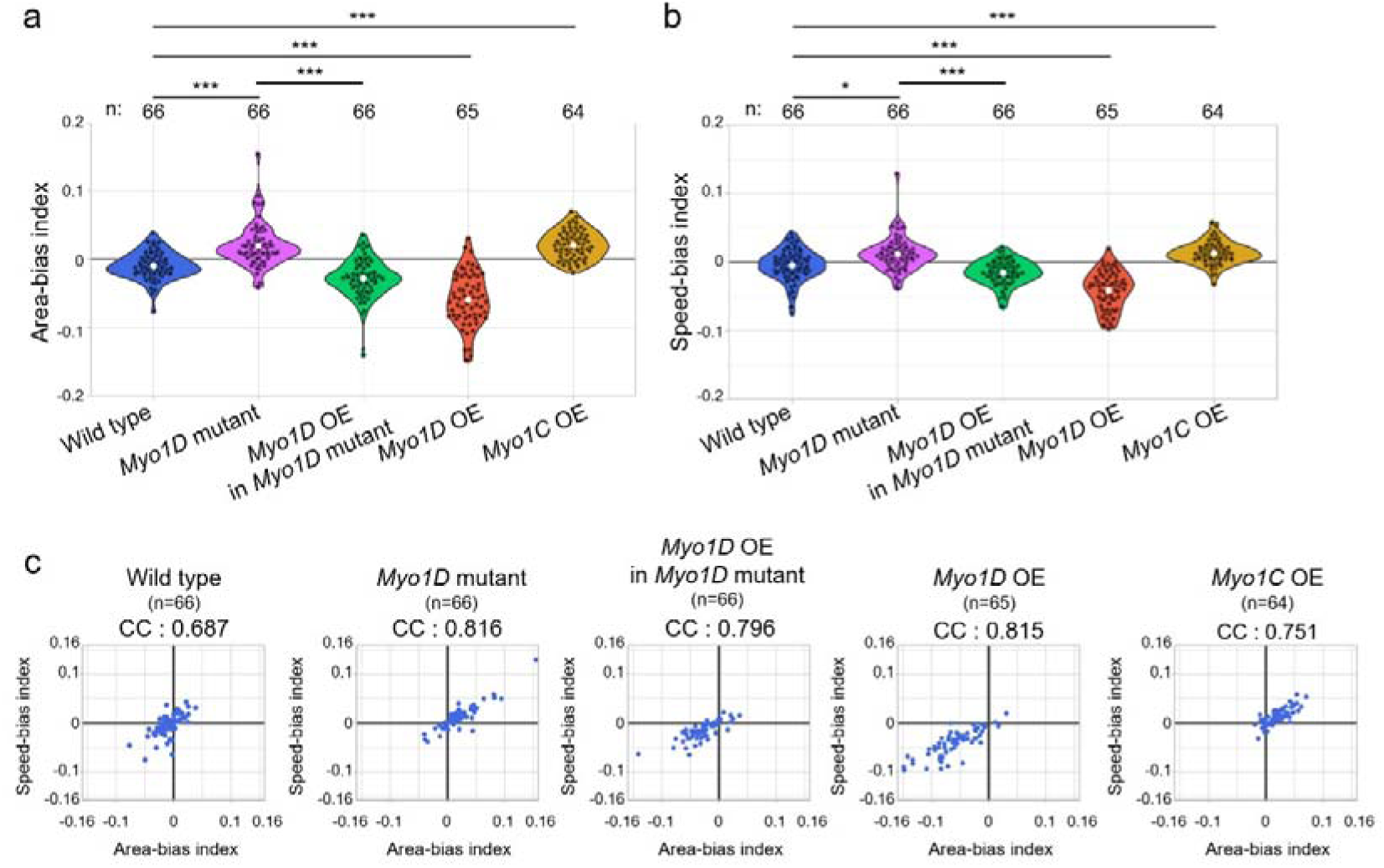
*Myo1D* and *Myo1C* expanded the intracellular area showing the CW and CCW rotations of F-actin, respectively. **a** and **b,** Violin plots showing the area index (a) and the speed index (b) of F-actin flow (plus and minus values represent CW and CCW biases, respectively) in macrophages with genotypes indicated at the bottom. White dots indicate the average value. The numbers of the analyzed macrophages are indicated at the top of the graphs. The *P*-values were obtained using Steel–Dwass; * and *** represent *P* < 0.05 and *P* < 0.0001, respectively. **c,** Scatter plot of the area index and the speed index of F-actin flow in the macrophages of genotypes indicated at the tops. CC represents the correlation coefficient between the area and speed indices calculated using Spearman’s rank correlation coefficient. The numbers of the examined macrophages are shown in parentheses.

We speculated that the CW or CCW bias of chiral F-actin domains can be achieved via two potential mechanisms. First, *Myo1D* and *Myo1C* may preferentially expand the chiral F-actin domains of CW and CCW, respectively. Second, *Myo1D* and *Myo1C* may preferentially accelerate the speed of CW and CCW F-actin flow, respectively. To test the first possibility, we defined the areas of the CW- and CCW-chiral F-actin domains as the total numbers of pixels corresponding to the CW- and CCW-chiral vectors, respectively (Table 3). The mean values of these areas showed significant CW or CCW bias corresponding to the respective bias of the chirality index of F-actin flow in all tested genotypes of macrophages; for example, significant CW-bias in the area of the chiral F-actin domain was observed in the wild type (Table3). To quantitatively evaluate the effects of *Myo1D* and *Myo1C* on the areas of the CW or CCW chiral F-actin domains, we determined the index representing the dominance of the area occupied by CW- or CCW-chiral F-actin domains as “area-bias index” (CCW pixel number − CW pixel number) / (CCW pixel number + CW pixel number). We found that the area-bias index of *Myo1D* mutant macrophages significantly shifted to positive (CCW-dominant expansion) values compared with that of the wild type (Fig.3a). This shift was converted to negative (CW-dominant expansion) values by the *Myo1D* overexpression in these mutant macrophages (Fig. 3a). Furthermore, the overexpression of *Myo1D* and *Myo1C* resulted in significant biases to negative and positive values of the area-bias index, respectively, as compared with that of the wild type. Thus, the CW and CCW biases of the area-bias index in various genetic conditions were parallel to those of the chirality index of F-actin flow (Fig. 1d, 3a).

**Table 3.** Average number of pixels demonstrating CW or CCW chiral F-actin flow. The *P*-values of the statistical significance of CW or CCW differences in the average pixel numbers demonstrating the CW or CCW chiral vector of the F-actin flow in the macrophages of the indicated genotypes were calculated via two-tailed one-sample t-test. *** represents *P* < 0.0001.

| Genotype | Average of pixels of CCW | Average of pixels of CW | p-value |
| --- | --- | --- | --- |
| Wild type | 91765.56 | 93961.34 | *** |
| *Myo1D* mutant | 109218.55 | 106081.94 | *** |
| *Myo1D* OE in *Myo1D* mutant | 100260.93 | 105861.41 | *** |
| *Myo1D* OE | 107536.06 | 121356.24 | *** |
| *Myo1C* OE | 103977.13 | 99574.86 | *** |

To evaluate the second possibility, we measured the effects of *Myo1D* and *Myo1C* on the acceleration of F-actin flow. We calculated the mean speeds represented by all F-actin chiral vectors with respective CW and CCW directionalities in each macrophage. We also obtained the averages of these speeds from macrophages with the genotypes tested (Table 4). We found that the bias in the average speeds with CW and CCW directionalities was not statistically significant in wild-type macrophages (Table 4); by contrast, they demonstrated a significant CW bias in the chiral F-actin domains (Table 3). Therefore, the selective area expansion of the chiral F-actin domains might have a more profound effect on determining the chirality index of F-actin flow rather than the CW or CCW-biased acceleration of F-actin flow. However, except for the wild type, the macrophages of all other tested genotypes showed significant CW- and CCW-biases of the average speeds, which were parallel to those of the chirality index of the F-actin flow (Table 4 and Fig. 1d). To quantitatively analyze the effects of *Myo1D* and *Myo1C* on the CW or CCW speeds of F-actin flow, we obtained a “speed-bias index” (average CCW speed − average CW speed / average CCW speed + average CW speed). We found that these effects were similar to those of the chirality index of F-actin in all the tested genetic conditions (Fig. 3b). Our analysis of the correlation coefficients between the area-bias and speed-bias indices revealed that they were strongly correlated in all examined genetic conditions (Fig. 3c). Considering the statistical significance observed in the CW-bias in the area of the chiral F-actin domain but not in the average speeds of F-actin flow in wild-type macrophages, our results suggested that the areal expansion of the chiral F-actin domains might have a more profound effect on determining the chirality index of F-actin flow than the CW- or CCW-biased acceleration of F-actin flow (Table 3 and 4). However, as the area-bias and speed-bias indices were strongly correlated (Fig. 3c), these two factors mutually reinforced each other. For example, the expansion of CW or CCW chiral F-actin domain might speed up respective chiral F-actin flow through the orchestration of mechanical force.

**Table 4.** Average speed of CW or CCW chiral F-actin flow. The *P*-values of the statistical significance of CW or CCW differences in the average size of the chiral vector of CW or CCW F-actin flow in the macrophages of the indicated genotypes were calculated using a two-tailed one-sample t-test. **, *P* < 0.001; ***, *P* < 0.0001.

| Genotype | Average speed of CCW actin flow | Average speed of CW actin flow | P-value |
| --- | --- | --- | --- |
| Wild type | 1.15 | 1.17 | 0.08 |
| *Myo1D* mutant | 1.10 | 1.07 | ** |
| *Myo1D* OE in *Myo1D* mutant | 1.17 | 1.21 | *** |
| *Myo1D* OE | 1.50 | 1.63 | *** |
| *Myo1C* OE | 1.63 | 1.59 | *** |

### Myo2 is essential for Myo1D and Myo1C to direct the chiral F-actin flow

In general, Myosin2 (Myo2) plays a major role in generating mechanical force that moves F-actin^27,28^. Furthermore, we previously revealed that *Myo1D* depends on Myo2 to induce dextral cell chirality in hindgut epithelial cells and the dextral rotation of the hindgut^29^. Therefore, in the present study, we examined whether Myo2 helps induce the chiral F-actin flow in macrophages. Myo2 activity is suppressed by the knockdown against *spaghetti squash* (*sqh*), encoding the regulatory light chain of Myo2 mediated by RNA interference (RNAi)^30,31^. RNAi against *sqh* was performed by expressing *UAS-sqh-RNAi* in wild-type macrophages and in macrophages overexpressing *Myo1D* or *Myo1C*; their chirality index of F-actin flow was calculated (Fig. 4a). We found that the CW or CCW-bias of the chirality index disappeared upon RNAi against *sqh* under these three conditions; conversely, RNAi against *Luciferase* (control) did not affect the CW or CCW-bias (Fig. 4a and Table 5). Therefore, *Myo1D* and *Myo1C* required a Myo2 activity to induce the CW and CCW chiral F-actin flows, respectively. To reveal the potential relevance between the chiral dynamics of F-actin and Myo2, we also simultaneously analyzed the chiral flow of Myo2 (visualized by *squ* tagged with *GFP* and expressed under the control of its promoter)^32^ and F-actin in macrophages. The chirality index of Myo2 flow showed a significant CW bias in wild-type and *Myo1D*-overexpressing macrophages and CCW bias in *Myo1C*-overexpressing macrophages, which are similar to the CW or CCW biases observed in the chirality index of F-actin flow in these macrophages (Extended Data Fig. 3a, b and Table 6). In these cells, we obtained the correlation coefficients between the chirality indices of F-actin and Myo2 flows (Fig. 4b). We found that the correlation coefficients between these two indices were 0.0791, 0.535, and 0.0115 in wild-type and *Myo1D*- or *Myo1C*-overexpressing macrophages, respectively (Fig. 4b). Therefore, in each macrophage with the *Myo1D* overexpression, the extent of CW or CCW biases of F-actin and Myo2 flows coincided with each other; conversely, such concordance was not observed in wild-type and *Myo1C*-overexpressing macrophages. Therefore, although Myo1D and Myo1C depend on Myo2 to induce the chiral F-actin flows, the underlying mechanisms may vary.

**Fig. 4.**
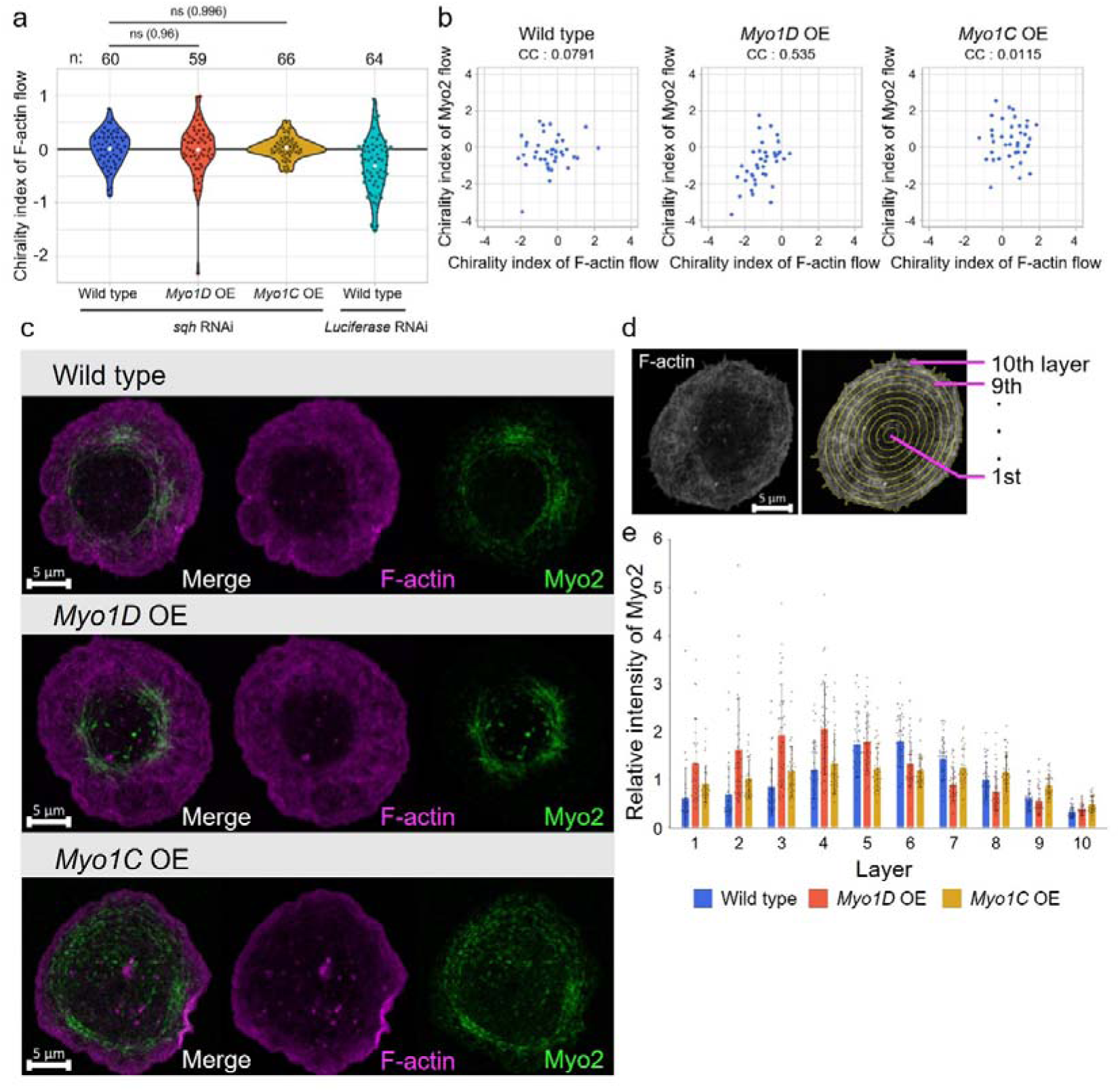
*Myo2* is essential for the dextral and sinistral chirality of the F-actin flow induced by *Myo1D* and *Myo1C*, respectively. **a,** Violin plots showing the chirality index of F-actin flow (plus and minus values represent CCW and CW biases, respectively) in the macrophages of the genotypes indicated at the bottom. These macrophages were treated with RNAi against *sqh* (encoding the regulatory light chain Myo2) or *Luciferase* (control). White dots indicate the average value. The numbers of the analyzed macrophages are indicated at the top of the graphs. The *P*-values were obtained using Steel–Dwass, and ns represents *P* > 0.05. **b,** Scatter plot of the chirality indices of F-actin flow and Myo2 flow in the macrophages of the genotypes indicated at the tops. CC represents the correlation coefficient between the chiral indices of Myo2 flow and actin flow calculated by using Spearman’s rank correlation coefficient. **c,** Typical example of the intracellular distribution of Lifeact-mCherry (F-actin, magenta) and sqh-GFP (Myo2, green) in the macrophages of the indicated genotypes. The left panels are the merged images of the middle and right panels. **d,** The procedure for defining the intracellular layers to quantify the distribution of Myo2. Using the fluorescent signal of F-actin, the cell contour is approximated as an ellipse, and the intracellular region is divided into 10 layers with concentric ellipses (yellow lines). **e,** Bar graphs showing the relative intensity of Myo2 in the 10 layers. Blue, orange, and yellow bars represent wild-type (31 cells), *Myo1D*-overexpressing (*Myo1D* OE, 38 cells), and *Myo1C*-overexpressing (*Myo1C* OE, 35 cells) macrophages. The error bar shows the standard error. Scale bars in c and d represent 5 μm.

**Table 5.** Statistical significance of biases in CW or CCW chiral F-actin flow in macrophages, where Myo2 activity was suppressed by RNAi (*sqh* RNAi). The *P*-values of the statistical significance of CW or CCW biases in the chiral vector of F-actin flow in the macrophages of the indicated genotypes and treated with *sqh* RNAi were calculated via a one-sample one-sided Wilcoxon signed-rank test.

| Genotype | *sqh* RNAi<br>in wild type | *sqh* RNAi<br>in *Myo1D* OE | *sqh* RNAi<br>in *Myo1C* OE | *Luciferase* RNAi<br>in wild type |
| --- | --- | --- | --- | --- |
| ***P* value** | **0.716** | **0.637** | **0.205** | **$8.932 \times 10^{-6}$** |

**Table 6.** Statistical significance of biases in CW or CCW chiral F-actin and Myo2 flow in macrophages, where F-actin and Myo2 were simultaneously labeled. The *P*-values of the statistical significance of CW or CCW biases in the chiral vector of F-actin or Myo2 flow in the macrophages of the indicated genotypes were calculated via a one-sample one-sided Wilcoxon signed-rank test.

| Chiral index of actin flow |  |  |  |
| --- | --- | --- | --- |
| Genotype | Wild type | *Myo1D* OE | *Myo1C* OE |
| *P* value | 0.00152 | $2.001 \times 10^{-10}$ | 0.00124 |
| Chiral index of *sqh* flow |  |  |  |
| *P* value | 0.0331 | $1.321 \times 10^{-5}$ | 0.0263 |

### *Myo1D* and *Myo1C* distinctively affect Myo2 distribution

To reveal the functional relationship between two class I myosins and Myo2, we analyzed the effects of *Myo1D* and *Myo1C* on the subcellular localization of Myo2. We detected Myo2 visualized by *sqh-GFP* and F-actin labeled by overexpressing *UAS-Lifeact-mCherry* in wild-type macrophages and macrophages overexpressing *Myo1D* or *Myo1C* (Fig. 4c, Extended Data Fig. 4a-c). Myo2 is generally enriched in transverse arcs, where F-actin assembles with antiparallel polarities in various species^33,34^. To analyze the intracellular distribution of Myo2 quantitatively, we measured the relative fluorescent intensities of Myo2 along the distance from the cell center (Fig. 4d and e). We divided the intracellular areas in each macrophage into 10 layers from the center (layer 1) to the margin (layer 10) of each cell (Fig. 4d). We analyzed the relative intensities representing Myo2 in each layer (Fig. 4e). In wild-type macrophages, the intensity of Myo2 peaked at the 5–6 layers (Fig. 4c and e). We found that the *Myo1D* overexpression shifted the distribution of Myo2 to the inner (1–4) layer compared with that of the wild type (Fig. 4c and e). However, the *Myo1C* overexpression resulted in a more uniform distribution of Myo2 (Fig. 4e). Therefore, *Myo1D* and *Myo1C* distinctively affected the intracellular distribution of Myo2.

### Myo1D induces the self-organization of F-actin ring with chiral rotation in the *in vitro* motility assay

Myo1D propels each F-actin CW when viewed from the myosin side (from the bottom of a glass slide) in an *in vitro* motility assay^16,35^. However, with previous results, the chiral motion of individual F-actin induced by Myo1D *in vitro* cannot be easily connected to the CW bias of chiral F-actin flow observed at the cellular level in *Myo1D*-overexpressing macrophages because the chiral motion of each F-actin merely induces multiple and local CW rotations on the cytoplasmic side of the plasma membrane in a previous *in vitro* motility assay^16^. Recently, we discovered that *Chara corallina* myosin XI (*Cc*XI) also induces a similar CW F-actin movement in an *in vitro* motility assay^36^. Furthermore, we modified the *in vitro* motility assay through which the F-actin concentration was adjusted to near-physiological concentrations; we found that *Cc*XI self-organized F-actin into ring□shaped F-actin bundles rotating CW, designated as “actin chiral ring (ACR)” ^36^(Fig. 5a). Thus, we performed the modified *in vitro* motility assay by using Myo1D and Myo1C to check whether they could also form the ACRs (Fig. 5a). We revealed that Myo1D induced the ACRs rotating CW (Fig. 5b). This CW rotation of the ACR implied that Myo1D has an activity to bundle F-actin with the same barbed pointed end polarity because F-actin should face their pointed ends to the CW direction as the ACR always rotates CW; subsequently, parallel and annular polarities are introduced into the ACR (Fig. 5b). The mean diameter of these ACRs is 5.4 ± 0.5 µm, which is similar to the diameter of macrophages (Fig. 1b). Therefore, we speculated that a cell-sized flow of F-actin bundles with parallel and annular polarities may be induced in *Myo1D*-overexpressing macrophages. Conversely, when a modified *in vitro* motility assay was performed with Myo1C, an amorphous flow with local and random directionality was observed (Fig. 5c). This result suggested that the CCW chirality of the Myo1C-induced F-actin flow in macrophages could not be solely explained by the Myo1C activity in the modified *in vitro* motility assay.

**Fig. 5:**
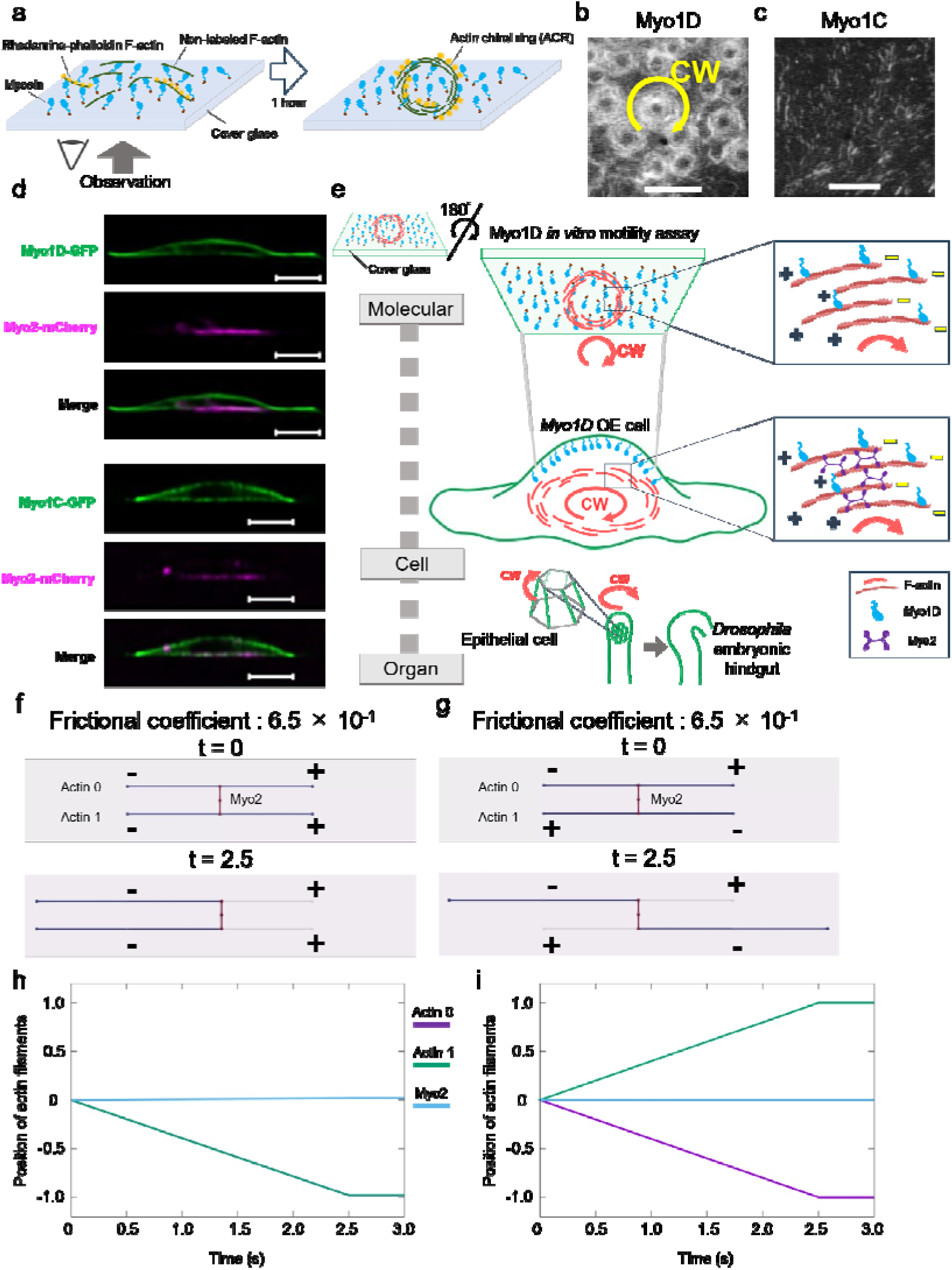
Myo1D formed the F-actin chiral ring rotating CW in a modified *in vitro* motility assay. **a,** A diagram showing the modified *in vitro* motility assay. **b** and **c,** Snapshots of fluorescent-labeled F-actin in the modified *in vitro* motility assay involving Myo1D (b) and Myo1C (c). Myo1D induced the F-actin chiral ring rotating CW (yellow rotation arrow), and Myo1C induced an amorphous flow with random directionality. Scale bars represent 10 μm. **d,** Vertical sections of macrophages under a Lattice Lightsheet microscope. The upper and bottom panels show the typical examples of the macrophages expressing *sqh-mCherry* (Myo2) (magenta) and *UAS-Myo1D-GFP* or *UAS-Myo1C-GFP* (green). Scale bars represent 5 μm. **e,** A diagram showing a potential mechanism of dextral cell and organ chirality formation. As revealed in *the in vitro* motility assay, Myo1D annularly and parallelly polarized F-actin fibers. In macrophages, Myo1D is mostly localized to the dorsal (upper) plasma membrane, which imitates the *in vitro* motility assay placed upside down and potentially introduces annularly and parallelly polarized F-actin fibers in the cytoplasm. In the cytoplasm, the displacement of Myo2 toward the barbed end of the annularly polarized F-actin can drive the LR-directional rotation of F-actin, which accounts for the CW flow of F-actin in macrophages once Myo1D is overexpressed. Subsequently, the CW flow of F-actin induces the cell twisting of the epithelial cells of the hindgut, which consequently drives the dextral rotation of the hindgut tube^1^. **f** and **g,** Simulations of translocation of two actin filaments under the condition of friction coefficient for Myo2 (6.5 × 10^−1^). Actin filaments are parallel (f) and antiparallel (g). t indicates the time point. Symbol – and + mean minus (pointed) and plus (barbed) end of actin filaments, respectively. **h** and **i**, The graphs represent the positional changes of actin filaments over time. h is the result of simulation (f) and i is the result of simulation (g). Blue, green, and magenta lines indicate the positional changes of Myo2 and each of the two actin filaments, respectively. Magenta line (Actin 0) and green line (Actin 1) overlap (h).

To examine the potential link between the ACR in the *in vitro* motility assay and the chiral F-actin flow in macrophages, we observed the three-dimensional intracellular localization of Myo1D and Myo1C in macrophages by using a Lattice Lightsheet microscope. As found in Z-section images, Myo1D and Myo1C were enriched in the dorsal (upper) plasma membrane (Fig. 5d). Thus, we presumed that the cytoplasmic surface of the dorsal plasma membrane may correspond to the glass surface, where Myosin Is were attached in our *in vitro* motility assay (Fig. 5e). In this case, Myo1D may introduce a similar annular polarity with the barbed pointed ends into the cytoplasmic F-actin in macrophages; consequently, it may allow Myo2 to propel the cytoplasmic F-actin along their polarity formed by Myo1D. This phenomenon explains the CW rotation of F-actin flow upon the *Myo1D* overexpression. To confirm this idea, we performed mathematical simulations in which a Myo2 filament drives two F-actin filaments with parallel or antiparallel polarity (Extended Data Fig. 5). In these simulations, we assumed that a single Myo2 filament and two actin filaments were anchored in the cytoplasm, subject to frictional forces from the surrounding cytosol. In the initial configuration, the two actin filaments were aligned with parallel polarity. Under this arrangement, both filaments moved toward the pointed (minus) end when the friction coefficient of Myo2 was higher than that of F-actin (Fig. 5f and 5h). However, this movement was not observed when Myo2 had a lower friction coefficient than F-actin (Extended Data Fig. 6a and 6c). The relationship between the displacement of actin filaments at the end of the simulations and the ratio of friction coefficients (Myo2/actin filaments) revealed that a higher friction coefficient of Myo2 caused the movements of the actin filaments toward the pointed (minus) end (Extended Data Fig. 6e). Moreover, if the polarities of the two actin filaments were not aligned, they did not move in the same direction (Fig. 5g, 5i, Extended Data Fig. 6b and 6d). Therefore, our simulations support that Myo2 propels the parallelly polarized F-actin toward the pointed end. On the basis of these results, we proposed a model with an annular and parallel polarity of the barbed pointed end into the ring of cytoplasmic F-actin bundles, which are introduced by Myo1D; the model may be circled through the Myo2 activity and drive the CW rotational motion of F-actin in the cytoplasm (Fig. 5e)

## Discussion

In *Drosophila*, the enantiomorphic states of the body, organ, and cell chirality, including those induced *de novo*, are determined by *Myo1D* and *Myo1C in vivo*^1,13–15^. To reveal the underlying mechanisms responsible for the opposing activities of *Myo1D* and *Myo1C* in directing the enantiomorphic states of chirality, various properties of Myo1D and Myo1C were compared, including their ATPase kinetics, the duration of their membrane retention, and gliding directionality and velocity of F-actin in the *in vitro* motility assay^16,35^. However, none of them explains the opposing activities of Myo1D and Myo1C in cell chirality formation. In this study, we revealed that *Myo1D* and *Myo1C* dictate the enantiomorphic (CW and CCW) states of F-actin flow in cells; this finding could explain a part of such underlying mechanisms. The CW and CCW biases of F-actin flow were reinforced by expanding the areas where F-actin rotates CW and CCW, respectively. Thus, *Myo1D* and *Myo1C* may affect the configuration and structure of the actin cytoskeleton, such as the polarity along the barbed pointed ends of F-actin as discussed below. Thus, the determination of the CW or CCW biases could be prearranged.

We also revealed that the activities of *Myo1D* and *Myo1C* to define the CW/CCW states thoroughly depended on Myo2. This observation agrees with our previous finding that the activity of *Myo1D* drives the dextral rotation of the hindgut requiring Myo2 ^29^. Thus, we speculated that mechanical forces rotating the cytoplasmic F-actin CW and CCW might be driven by Myo2 based on the polarities of F-actin bundles generated through *Myo1D* and *Myo1C*, respectively. Because the underlying mechanism by which polarity could be introduced into F-actin bundles, we revealed that Myo1D induces the formation of the ACR that rotates CW when viewed from the myosin side in the modified *in vitro* motility assay through which F-actin was added to a concentration equivalent to that of *in vivo*^36^. The CW rotation *in vitro* implies that the F-actin bundles forming the ACR composed of annularly and parallelly polarized F-actin concerning the barbed pointed ends because MyoID propels F-actin only toward its pointed end (minus end; Fig. 5e). In this study, we showed that Myo1D and Myo1C preferentially localize to the plasma membrane, more predominantly to the dorsal plasma membrane, in macrophages (Fig. 5d). Given that the specific association of *Myo1D* with the dorsal plasma membrane recapitulated the glass surface where Myo1D was attached in the modified *in vitro* motility assay, our model predicted the CW rotation of the ACR in the Myo1D-overexpressing macrophages (Fig. 5d and 5e). Hence, if Myo2 moves more slowly than F-actin does because of viscous resistance or anchoring, we proposed that the displacement of Myo2 toward the barbed end of the annularly polarized F-actin could propel it toward the pointed end and drive the CW flow of F-actin in macrophages overexpressing *Myo1D* (Fig. 5e). Our computational simulations demonstrated that Myo2 with a higher friction coefficient could more efficiently drive parallelly polarized F-actin filaments toward their pointed ends, supporting this hypothesis. However, an absolute polarization of F-actin is not required *in vivo*, and an inclination of polarity in the annulus of F-actin is probably enough to induce the chiral flow of F-actin.

Myo2 is associated with the transverse arcs of F-actin assembled with antiparallel polarities, leading to the enrichment of Myo2 in a ring□shaped or crescent structure^33,34^. Our studies revealed that the *Myo1D* and *Myo1C* overexpression interiorly and exteriorly changes the intracellular location where Myo2 is enriched, respectively, suggesting that the activities of *Myo1D* and *Myo1C* affect the transverse arcs of F-actin in macrophages. We speculated that parallelly and annularly polarized F-actin induced by *Myo1D* might suppress the formation of the transverse arcs of F-actin fibers in the outer region of the cytoplasm; consequently, the Myo2-enriched region could be moved interiorly.

In our *in vitro* motility assay, Myo1C propelled F-actin as an amorphous flow with local and random directionality (Fig. 5c). We speculated that this local flow of F-actin might also assemble F-actin in a manner with parallel polarity; however, unlike Myo1D, Myo1C did not induce the annularly polarized F-actin bundles. Therefore, Myo1C might place F-actin bundles with parallel polarity in an intracellular configuration that differed from that of Myo1D. Such distinct configurations might explain the difference in the expansion and scattering of transverse arcs of F-actin fibers with Myo2 enrichment between *Myo1D*- and *Myo1C*-overexpressing macrophages. These results also suggested that Myo1D and Myo1C induce the respective chiral F-actin flow through different mechanisms originating from the distinct structures of the polarized F-actin bundle; this result is consistent with our previous genetic observations that the sinistral activity of *Myo1C* is independent of *Myo1D*^14^.

## Methods

### Fly stocks

The following genes were used: *UAS-Lifeact-mGFP6* and *UAS-Lifeact-mCherry* (gift from Dr. Shinya Yamamoto, Baylor College of Medicine), encoding F-actin markers; *UAS-GFP-spd2* (gift from Dr. Yuu Kimata, ShanghaiTech University), encoding centrosome markers; *UAS-Redstinger*^37^ and *UAS-stinger*^38^, encoding nucleus markers; *UAS-Myo31DF-HA*^39^ and UAS-Myo1D-mGFP6 (in this study), encoding Myo1D; *UAS-GFP-*α*tub* (#BDSC7373), encoding GFP-tagged α-tublin; *UAS-Myo61F-HA*^39^and UAS-Myo1C-mGFP6 (in this study), encoding Myo1C; *UAS-Cnn-GFP* ^26^, encoding a centrosome marker; and *sqh-GFP* (BDSC#57145), encoding GFP-tagged *sqh*; *UAS-sqh-RNAi* (#BDSC33892), encoding a double-stranded RNA of *sqh*; *He-GAL4.Z* (#BDSC8699) was used as a GAL4 driver to induce the macrophage-specific expression^40^; *Myo31DF^L1^*^52^, an amorphic allele of *Drosophila Myo1D*^13^. All genetic crosses were performed on a standard *Drosophila* culture medium at 25 °C.

### Generation of UAS-Myo1D-mGFP6 and UAS-Myo1C-mGFP6

pUASt-Myo1D-Halotag was used as a template to construct UAS-Myo1D-mGFP6^41^. The *Kpn*I and *Eco*RI fragments, including a cDNA of Myo1D tagged with a HaloTag, were ligated into the *Kpn*I and *Eco*RI sites of the pBluescript II sk(−) plasmid. The construct was used as the template, and a Myo1D fragment with a linker (5′-TCCTCGAGCTCATCG) added to the 3′-end was PCR-amplified. An mGFP6 fragment was also PCR-amplified using UAS-Lifeact-mGFP6 as a template with a linker (5′-TCCTCGAGCTCATCG) and a *Not*I site attached to 5′- and 3′-ends, respectively. These two PCR fragments were combined via PCR, and the combined fragment was digested using *Eco*O109I and *Kpn*I. The resulting fragment was ligated into the *Eco*O109I and *Kpn*I sites of pBluscript-Myo1D-Halotag. Consequently, pBluescript-Myo1D-mGFP6 was obtained and digested with *Eco*RI and *Not*I. The EcoRI and NotI fragments were purified and cloned into the same restriction sites of pUASt to obtain pUASt-Myo1D-mGFP6.

pUASt-Myo1C-Halotag^41^ was used as a template to construct UAS-Myo1C-mGFP6, and pBluescript-Myo1C-Halotag was prepared in the same procedures to pBluescript-Myo1D-Halotag. A cDNA of Myo1C was PCR-amplified using Bluescript-Myo1C-Halotag as a template, but the overlapping region of pBluescript was added at the 5′-end, and the linker and mGFP6-overlapping region were attached at the 3′-end. A vector backbone comprising *mGFP6* was obtained via inverse PCR using pBluescript-Myo1D-mGFP6 as a template. The cDNA fragment of *Myo1C* and the vector backbone were combined through homologous recombination; consequently, pBluescript-Myo1C-mGFP6 was obtained. This construct was digested by *Eco*RI and *Not*I, and the fragment of Myo1C-mGFP6 was isolated. The fragment was cloned into the same restriction sites of pUASt to obtain pUASt-Myo1C-mGFP6.

The DNA sequences of *Myo1D* and *Myo1C* cDNAs and linkers were confirmed through DNA sequencing analysis. These constructs were integrated into the 68A4 P[*CaryP*]attP2 site of the third chromosome by using the *PhiC31*/attP/attB system to obtain the transgenic lines of pUASt-Myo1D-mGFP6 and pUASt-Myo1C-mGFP6^42^.

### Preparation of macrophages for live imaging

The genes of interest were specifically expressed in macrophages under the control of the UAS promoter driven by *He-GAL4.Z* by using the Gal4/UAS system. Body fluid containing these macrophages was obtained from the wound of the third instar larvae pricked by a glass needle and transferred onto Concanavalin A (Nakarai tesque)-coated cover glass in M3 insect media (Sigma) supplemented with 12.5% fetal bovine serum (BioWest), 0.25% polypeptide (Fujifilm), and 0.1% Tc Yeastolate (Thermofisher)^43^. In the experiments involving the inhibition of microtubule polymerization, colchicine (sigma) was added in the medium (final concentration: 10 μg/ml). The macrophages were cultured at 25 °C for 30 min, successively subjected to time-lapse imaging by using LSM 880, and processed via Airlyscan (Carl Zeiss) or Lattice Lightsheet microscopy (Carl Zeiss).

### Optical flow analysis of F-actin and Myo2 circumferential flows

F-actin and Myo2 were visualized using fluorescence derived from Lifeact and *sqh-GFP*, respectively, in cultured macrophages. Time-lapse fluorescent movies were obtained every 15 s for 10 min by using an LSM 880 confocal microscope (Carl Zeiss). In these time-lapse movies, the direction vector of optical flow representing all pixels was estimated through optical flow analysis by using OpenCV, Gunnar Farneback’s algorithm^22^.

The direction vector of optical flow was separated into two directions to obtain the chiral vector that represents the circumferential motion of F-actin and Myo2: one was toward the cell centroid, and the other was perpendicular to it (Fig. 1c). The latter represented the chiral vector with clockwise (minus) and counterclockwise (plus) directionalities. In this calculation, the cell center was defined as the centroid of the cell’s outline obtained by the ImageJ threshold function from the first image of time-lapse movies. The positive or negative values of chiral vectors representing all pixels were integrated into each cell to calculate the chirality index of F-actin and Myo2 flows.

### Visualization of chiral F-actin domains

Among the chiral vectors representing the movement of pixels for 10 min, one in every pixel along the X and Y axes was selectively shown in the first image of time-lapse movies. The starting points of the selected chiral vectors were placed at the corresponding pixels of the images and indicated by dots. These chiral vectors with CW and CCW directions were presented in yellow and magenta, respectively.

### Calculation of the area–bias index

All chiral vectors representing the pixel movement of 10 min were classified into two categories with CW or CCW directionality (CW- and CCW-chiral vectors). The total numbers of CW- and CCW-chiral vectors, which are proportional to the areas occupied by F-actin flow with CW and CCW chirality, respectively, were obtained in each macrophage. As an index representing the CW/CCW bias of the area occupied by CW and CCW chiral F-actin flow, the area–bias index was calculated using the following equation: (number of CCW-chiral vector − number of CW-chiral vector) / (number of CCW-chiral vector + number of CW-chiral vector).

### Calculation of the speed–bias index

The average magnitudes of the chiral vectors belonging to CW- or CCW-chiral vectors were calculated. The speed–bias index was calculated from the following formula to quantitatively analyze CW or CCW bias in the speed of the chiral F-actin flow: (number of CCW-chiral vector − number of CW-chiral vector) / (number of CCW-chiral vector + number of CW-chiral vector).

### Measurement of the rotation angle of the centrosome

*He-Gal4* simultaneously induced the misexpression of *UAS-Cnn-GFP* and *UAS-Redstinger* in macrophages to acquire time-lapse movies depicting centrosome rotation. The live images of macrophages cultured on Concanavalin A-coated cover glass were captured for 6 h (for the analysis of 30 min intervals) under a DeltaVison (GE) fluorescence microscope at 21 (±1) °C. The outline of the nucleus labeled by Redstinger was approximated as a circle fitted using the ImageJ threshold function in each time-lapse image. The center of the circle was computationally defined as the center of the nucleus by using the Fit as a circle function of ImageJ. At the same image, the position of the centrosome was manually assigned using the Fit as a circle function of ImageJ. Based on two-dimensional coordinates represented by the nuclear center and centrosome position, a line through two coordinates was formulated in each image. The angle between the lines formulated in two time-lapse frames (every 30 min for Fig. 2b) was calculated as angle *θ* with clockwise (minus) and counterclockwise (plus) directionalities.

### Analysis of correlation coefficients between the chirality index of F-actin and the centrosome rotation

*He-Gal4* simultaneously induced the expression of *UAS-Lifeact-mCherry* and *UAS-Cnn-GFP* in macrophages to acquire time-lapse movies depicting the F-actin flow and centrosome rotation simultaneously. Their live images were captured every 15 min for 20 min by using an LSM880 (Zeiss) confocal microscope. The chirality index of F-actin flow and the rotation angle of the centrosome were calculated as described above, and the correlation coefficients between them were obtained via Pearson analysis.

### Observation of the intracellular distribution of Myo1D and Myo1C

The three-dimensional images of macrophages were acquired using Lattice Lightsheet 7 (Carl Zeiss) at approximately 25 °C. The acquired images were deconvoluted using ZEN (Carl Zeiss) based on the “Nearest Neighbor” function. Subsequently, cross-sections were obtained from the pictures processed with 3D Crop (Fiji).

### Quantitative analysis of Myo2 and F-actin distributions in macrophages

A culture chamber with a silicon sheet (Extended Data Fig. 4a-c) was prepared to obtain higher-resolution images of Myo2 by using an upright microscope (LSM 880, Carl Zeiss). A silicon sheet with a cell culture space and a flow channel was placed on a glass slide and covered with a cover glass coated with Concanavalin A (Nakarai tesque). The suspension of macrophages expressing *sqh-GFP* and overexpressing *UAS-Lifeact-mCherry* by *He-Gal4* was poured into the flow channel, and they were allowed to adhere to the cover glass at 27 °C for 30 min. These macrophages were observed upside down via Airlyscan processing (Carl Zeiss). Based on the distribution of Lifeact-mCherry, the outlines of macrophages were defined using the function of elliptical approximation in Fiji. The major and minor axes of the ellipse were divided by 10, and the ellipse was sectioned into 10 layers, where the 10th layer faced the outline. The intensity of *sqh-GFP* and the area in each layer were measured, and the intensity was divided by the area of each layer to obtain the relative intensity of Myo2 per area unit in the layers in each macrophage. The relative intensity per area unit was divided by the relative intensity per area unit of the whole cell to standardize the relative intensity per area unit of the layers among the macrophages of each genotype.

### Preparation of Myo1C and Myo1D portions for *in vitro* motility assay

The. A *NcoI* site (first nucleotide) and an *AgeI site* (1035th and 1011th nucleotides in *Myo1C* and *Myo1D*, respectively) were added to the cDNAs of *Drosophila Myo1C* (UNIPROT ID: Q23979) and *Myo1D* (UNIPROT ID: Q23978) via PCR and then the fragments were inserted at the *NcoI* and *AgeI* sites of the pFastBac (Invitrogen, Carlsbad, CA, USA) CCM MD vector^44^. The resulting constructs, namely, pFastBacMyo1C and pFastBac Myo1D, encode Myo1C and Myo1D, respectively, with a FLAG-tag (DYKDDDDK) in the N-terminus and Myc-tag (EQKLISEEDL) and His8-tag (HHHHHHHH) in the C-terminus. Calmodulin is a light chain of the myosin I family^45^. An *XbaI* site (first nucleotide) and an *XhoI* site (149th nucleotide) were added to a cDNA of *Drosophila calmodulin* (UNIPROT ID: P62152) through PCR and inserted at the *BamHI* and *KpnI* sites of the pFastBac vector to create pFastBac CaM. pFastBacMyo1C, pFastBacMyo1D, and pFastBacCaM were transformed into DH10Bac to obtain the bacmids of baculovirus^46^. These bacmids were subsequently transfected into SF9 cells to produce baculoviruses carrying the *Myo1C*, *Myo1D*, and *CaM* constructs^46^.

Baculoviruses carrying the *Myo1C* or *Myo1D* construct were infected with baculoviruses carrying the *CaM* construct to High FiveTM cell culture (Invitrogen) to produce Myo1C and Myo1D proteins. The cells were harvested and washed in accordance with the standard procedure^46^. The pelleted cells were suspended in three-fold volume (ml) per cell weight (g) of buffer A (30 mM HEPES, pH 8.0, 200 mM NaCl, 50 mM KCl, 5 mM MgCl2, 10% glycerol, 10 mM β-mercaptoethanol, 0.5 µM *Chara corallina* calmodulin, and a mixture of protease inhibitors) containing 7 mM ATP. Three-fold volume (ml) per cell weight (g) of buffer A containing 1% Nonidet P-40 was added, and the ingredients were mixed. After incubation on ice for 15 min, the lysate was centrifuged at 228,000 × *g* at 4 °C for 30 min. The supernatant was mixed with 0.3 ml of nickel-nitrilotriacetic acid-agarose (Qiagen) in a 50 ml tube on a rotating wheel at 4 °C for 1 h. The resin suspension was then loaded on a column and washed with 30 ml of buffer A containing 30 mM imidazole and washed with 10 ml of buffer B (30 mM HEPES, pH 8.0, 50 mM KCl, 5 mM MgCl_2_, 30 mM imidazole, 10% glycerol, 10 mM *β*-mercaptoethanol, 0.5 µM *C. corallina* calmodulin, and a mixture of protease inhibitors) containing 2 mM ATP. Myo1C and Myo1D proteins were eluted with buffer B containing 250 mM imidazole. The eluted proteins were mixed with 0.3 ml of anti-FLAG M2 affinity resin (Sigma) in a 50 ml tube on a rotating wheel at 4 °C for 1 h. The resin suspension was loaded on a column, washed with 10 ml of buffer C (25 mM HEPES, pH 7.4, 150 mM KCl, 4 mM MgCl_2_, 10% glycerol, 0.5 mM DTT, 0.5 µM *C. corallina* calmodulin, and a mixture of protease inhibitors) containing 2 mM ATP, and washed with 30 ml of buffer D (25 mM HEPES, pH 7.4, 25 mM KCl, 4 mM MgCl_2_, 10% glycerol, 0.5 mM DTT, *C. corallina* calmodulin, and a mixture of protease inhibitors) containing 0.2 mM ATP. Myo1C and Myo1D proteins were eluted with buffer D containing 0.2 mg/ml of FLAG peptide (Sigma). Purified Myo1C and Myo1D proteins were rapidly frozen in liquid nitrogen and stored at −80 °C. *C. corallina* calmodulin (Q9LDQ9) was expressed in *Escherichia coli* strain BL21 (DE3) and purified via trichloroacetic acid method and affinity chromatography by using Phenyl Sepharose CL-4B^47^. G-actin was purified in accordance with a standardized protocol from acetone powder prepared from a rabbit skeletal muscle following a previously described method^48^.

### Modified *in vitro* motility assay using high-concentration actin

The *in vitro* motility assay was modified using actin concentrations closer to physiological levels than standard assays, as previously described^36^. Briefly, the purified Myo1C or Myo1D protein was immobilized on the coverslip by using an anti-c-Myc antibody ^44,49,50^. The assay chamber was subsequently infused with an assay buffer containing 0.005 mg/ml rhodamine-phalloidin-labeled actin and 0.1–0.3 mg/ml unlabeled actin without ATP. Rhodamine-phalloidin F-actin was pre-treated with Gelsolin (Sigma-Aldrich, product number: G8032) at a molar ratio of 1000:1 in the presence of 1 mM CaCl_2_; thus, its average length was approximately 5 µm although the average length of the actin filaments shortened because of severing events associated with their movement during the *in vitro* motility assay. After 10 min of incubation without ATP, the assay buffer with ATP (25 mM Hepes-KOH (pH 7.4), 25 mM KCl, 4 mM MgCl_2_, 10 mM ATP, 10 mM DTT, 30 µM *C. corallina* calmodulin, and an oxygen scavenging system (120 µg/ml glucose oxidase, 12.8 mM glucose, and 20 µg/ml catalase) was infused to remove unbound actin filaments. The time-lapse imaging of F-actin movement was conducted under a fluorescence microscope with a camera (WAT-910HX, Watec). Because the formation of the collective motion of actin filaments, including actin chiral rings, required 60–90 min, observations were conducted after incubation for 60–90 min.

### Measuring the diameter of the ACRs in the modified *in vitro* motility assay

The diameter of the ACRs were measured by the same method described previously^36^.

### Model description

We developed a mathematical model to demonstrate that a myosin filament can translocate actin filaments with aligned polarity. In this model, we consider two actin filaments (indexed by *i* = 0, 1) that undergo translational motion only; rotational motion is not considered (Extended Data Fig.5). A single myosin filament is assumed to be anchored to an object such as cytoplasmic structure. The myosin filament has two head domains at both ends, which are coarse-grained representations of multiple myosin heads. Each myosin head moves towards the plus end of the actin filament at a constant velocity *v*. We neglect the backward steps of the myosin heads due to thermal fluctuations because we focus on the averaged motion of the filaments. The inertia of the myosin filament is also negligible because of its sub-micrometer size, where viscous forces dominate. The following equations of motion were numerically integrated using the first-order Euler method. The simulation parameters are shown in Table 7.

**Table 7.** Simulation parameters.

| Parameter | Value(s) | Source/Note |
| --- | --- | --- |
| Actin length ($\mu\text{m}$) | 2.0 | |
| Initial distance between the actin filaments ($\mu\text{m}$) | $3.0 \times 10^{-1}$ | The length of non-muscle myosin bipolar filament<sup>51</sup>. This distance is set assuming that the bipolar myosin filament spans between the two actin filaments. |
| Actin friction coefficient, $\gamma$ (pN/($\mu\text{m/s}$)) | $6.5 \times 10^{-3}$ | Cylindrical object estimation<sup>52</sup>, assuming the actin filament diameter and cytoplasmic viscosity in the reference. |
| Myosin friction coefficient, $\Gamma$ (pN/($\mu\text{m/s}$)) | $6.5 \times 10^{-5}$, $1.0 \times 10^{-4}$, $3.0 \times 10^{-4}$, $6.5 \times 10^{-4}$, $1.0 \times 10^{-3}$, $3.0 \times 10^{-3}$, $6.5 \times 10^{-3}$, $1.0 \times 10^{-2}$, $3.0 \times 10^{-2}$, $6.5 \times 10^{-2}$, $1.0 \times 10^{-1}$, $3.0 \times 10^{-1}$, $6.5 \times 10^{-1}$, 1.0, 3.0, 6.5 | Values used to test the effect of anchor friction. |
| Myosin spring constant, $k$ (pN/$\mu\text{m}$) | $2.0 \times 10^3$ | <sup>53</sup> |
| Myosin speed, $v$ ($\mu\text{m/s}$) | $4.0 \times 10^{-1}$ | <sup>52</sup> |
| Time step (s) | $1.0 \times 10^{-9}$ | This small-time step ensures the numerical stability of the simulation. |
| Total simulation time (s) | 3.0 |  |

### Actin filament motion

The motion of actin filament *i* is described by its center-of-mass coordinate, 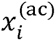, along the *x*-axis (which is parallel to the direction of the translational motion):

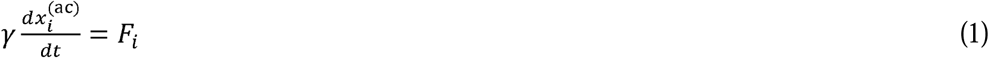

where *γ* is the friction coefficient of actin filament *i*, and *F* is the *x*-component of the force acting on actin filament *i*.

### Myosin head motion

The motion of the myosin head bound to actin filament *i* is described as follows:

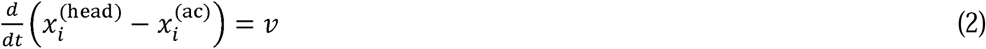

where *x*^⁽⁾^ is the coordinate of the myosin head bound to actin filament *i*, and *v* is the constant velocity of the myosin head along the *x*-axis.

### Force between myosin head and actin filament

The force exerted on actin filament *i* by the myosin head is given by:

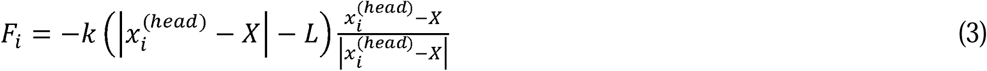

where *k* is the spring constant, *L* is the natural length (i.e., the stress-free distance) between the myosin head and the center of mass of the myosin filament (*X*), and *X* is the center-of-mass coordinate of the myosin filament. Since *L* is expected to be small compared to the overall filament displacements, we set *L*= 0 for simplicity.

### Myosin filament motion

The equation of motion for the center of mass of the myosin filament (*X*) is:

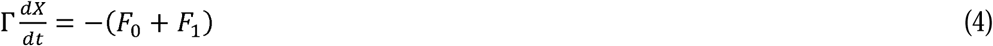

where *Γ* is the friction coefficient of the myosin filament, including the friction from the anchor object. *F*_0_ and *F*_l_ are the forces exerted by the myosin heads on the two actin filaments (*i*=0 and 1), respectively.

### Statistical analysis

Data were statistically analyzed using R (version 3. 6. 1). *P*-values were calculated via Steel–Dwass (Fig. 1d, 2d, 3a, 3b, 4a, Extended Data Fig. 3a, and 3b). One-sample Wilcoxon signed rank tests were performed (Tables 1, 2, 5, and 6). Pearson’s correlation coefficient was obtained (Fig. 2e). Spearman’s rank correlation coefficient was obtained (Fig. 3d and 4b).

## Acknowledgements

We thank the members of the Matsuno Laboratory for their valuable advice and discussions. We also thank Dr. Shinya Yamamoto, Dr. Yuu Kimata and the Bloomington *Drosophila* Stock Center (Indiana University) for *Drosophila* stocks, and Dr. Takahiro Fujiwara (ZEISS-iCeMS Innovation Core at the iCeMS Analysis Center, Kyoto University), Dr. Tatsushi Igaki (Kyoto University), Dr. Kiichiro Taniguchi (Kyoto University) for the observation with using Lattice Lightsheet microscopy. This work was supported by the Grant-in-Aid for Scientific Research on Innovative Areas (15H05863) to K.M.; by the Grant-in-Aid for Transformative Research Areas (24H01284) to K.M.; and by the Grant-in-Aid for JSPS Fellows (22J20544) to A.Y.

## Author contributions

K.M. and K.I. conceived and directed the study. A.Y., T.S., H.T., D.K., Y.A., and T.H. performed the observation of macrophages and data analysis. M.A. supervised mathematical calculations such as the chiral index, the rotation angle of movements of the centrosome, the speed-bias index, and the area-bias index. T.H., K.Y. and M.I. performed the modified *in vitro* motility assay. Y.I. developed the mathematical model and performed the simulation and the data analysis. Y.S. helped with the observation of cells using the Lattice Lightsheet microscope. K.M., K.I., A.Y., and F.L.N. wrote the draft. All authors approved the final version of the manuscript.

---

**Extended Data Fig. 1.**
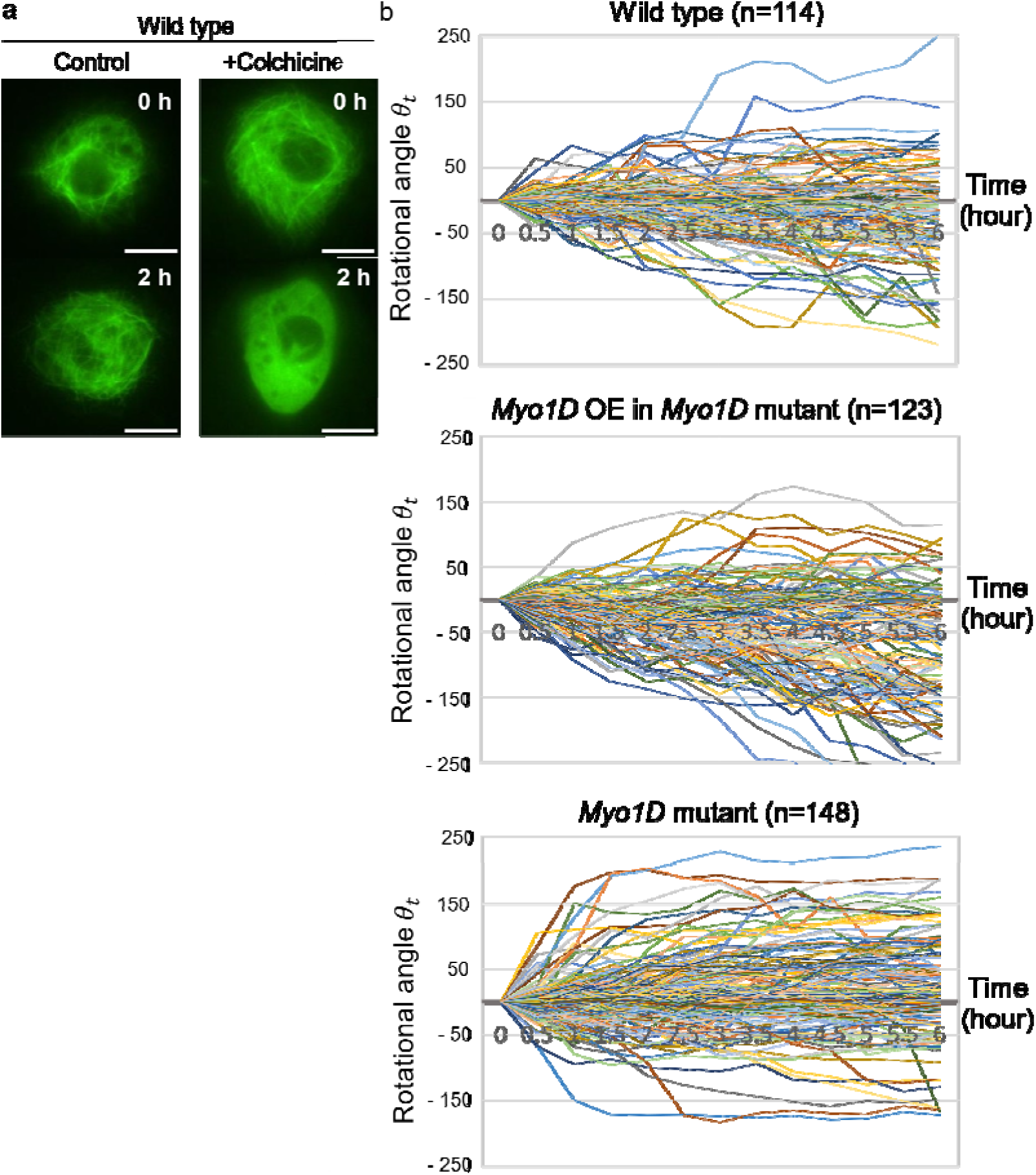
Rotation angles of the centrosome in wild-type and *Myo1D* mutant macrophages and *Myo1D* mutant macrophages overexpressing *Myo1D*. **a,** Time-lapse images of wild type macrophages expressing *GFP-*α*tub* to visualize microtubule structure. Left: wild type macrophage treated with water (control), and right: wild type macrophages treated with an inhibitor against polymerization of microtubules, colchicine. Each time point is indicated at the top right. Scale bars are 10 μm. **b,** Rotation angles (*θ_t_*) of the centrosome from t=0 min every 30 min (0.5 h) for six h (plus and minus values represent the CCW and CW direction of the rotation, respectively) of macrophages with genotypes indicated at the top. Polygonal lines shown in different colors represent the results of each macrophage. The genotypes of macrophages are shown at the top of each graph. The numbers of macrophages examined are shown in parentheses.

**Extended Data Fig. 2.**
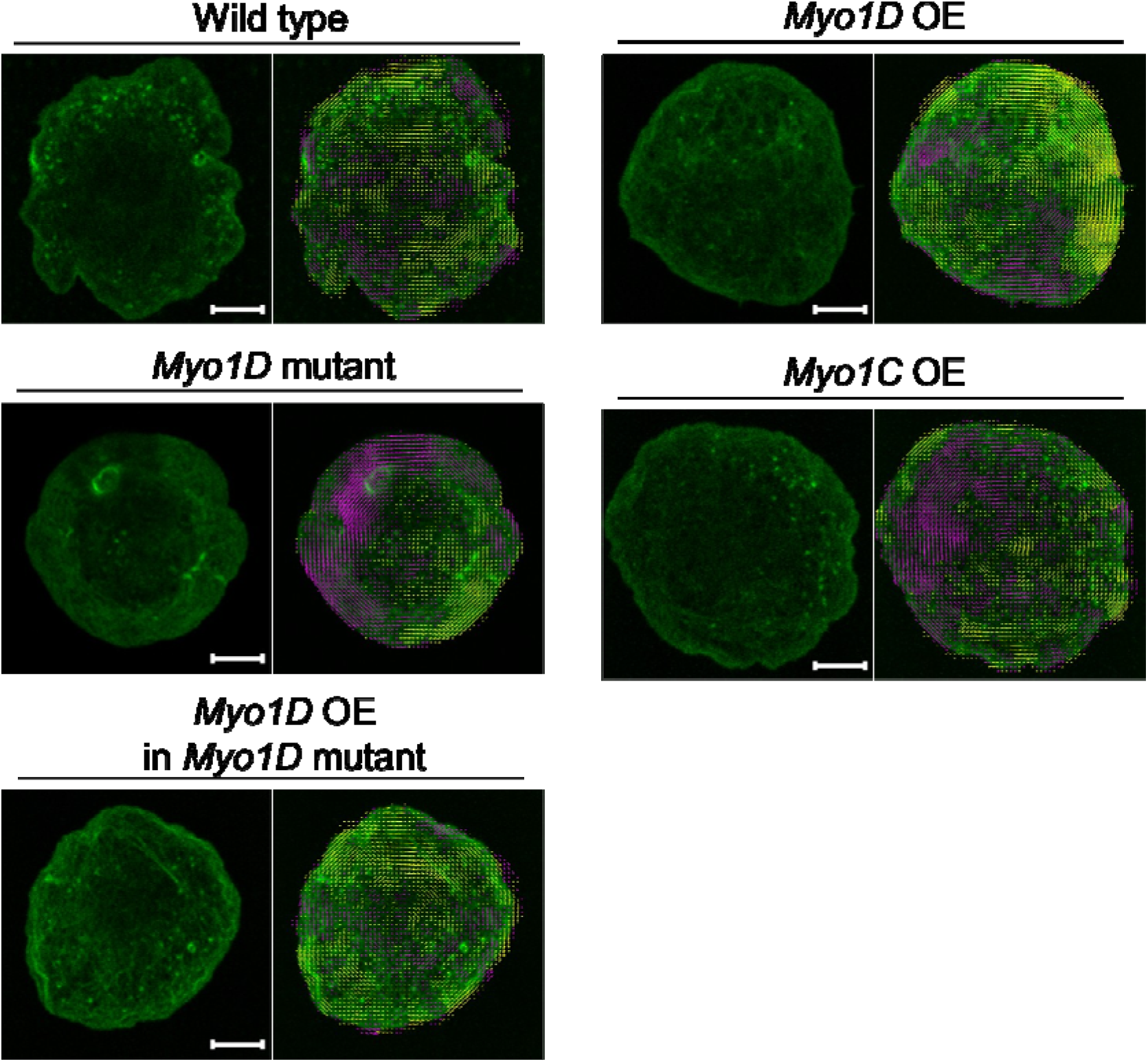
Typical images of the macrophages of the indicated genotypes overexpressing Lifeact-GFP (left) and chiral vectors drawn on the images (right). The F-actin chiral vectors are visualized for every 10 pixels with CW (yellow) and CCW (magenta). Scale bars represent 5 μm.

**Extended Data Fig. 3.**
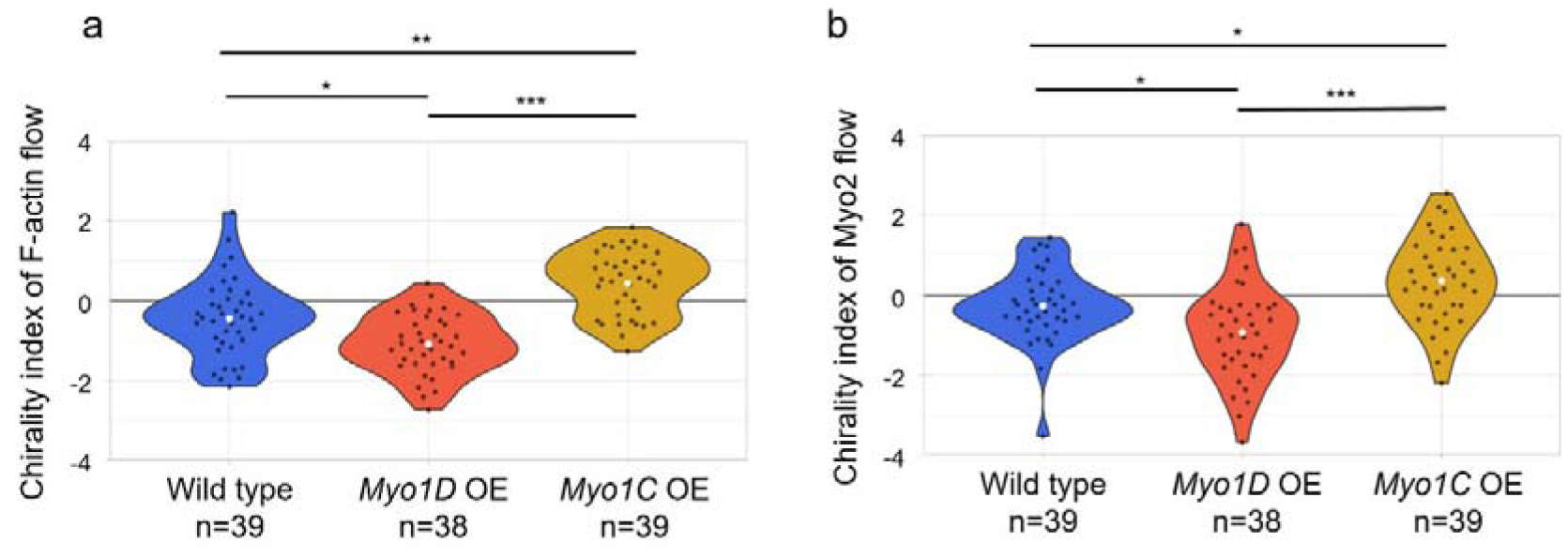
Myo1D and Myo1C induce CW and CCW rotation of not only actin flow but also Myo2 flow. **a** and **b**, Violin plots showing the chirality index of F-actin (a) and Myo2 (b) flows (plus and minus values represent CCW and CW biases in the direction of the flows, respectively) in macrophages with the genotypes indicated at the bottom. White dots indicate the average values. The numbers of macrophages analyzed are indicated at the bottom of the graphs. The *P*-values were obtained using Steel–Dwass, and *, **, and *** represent *P* < 0.05, *P*< 0.001, and *P* < 0.0001, respectively.

**Extended Data Fig. 4.**
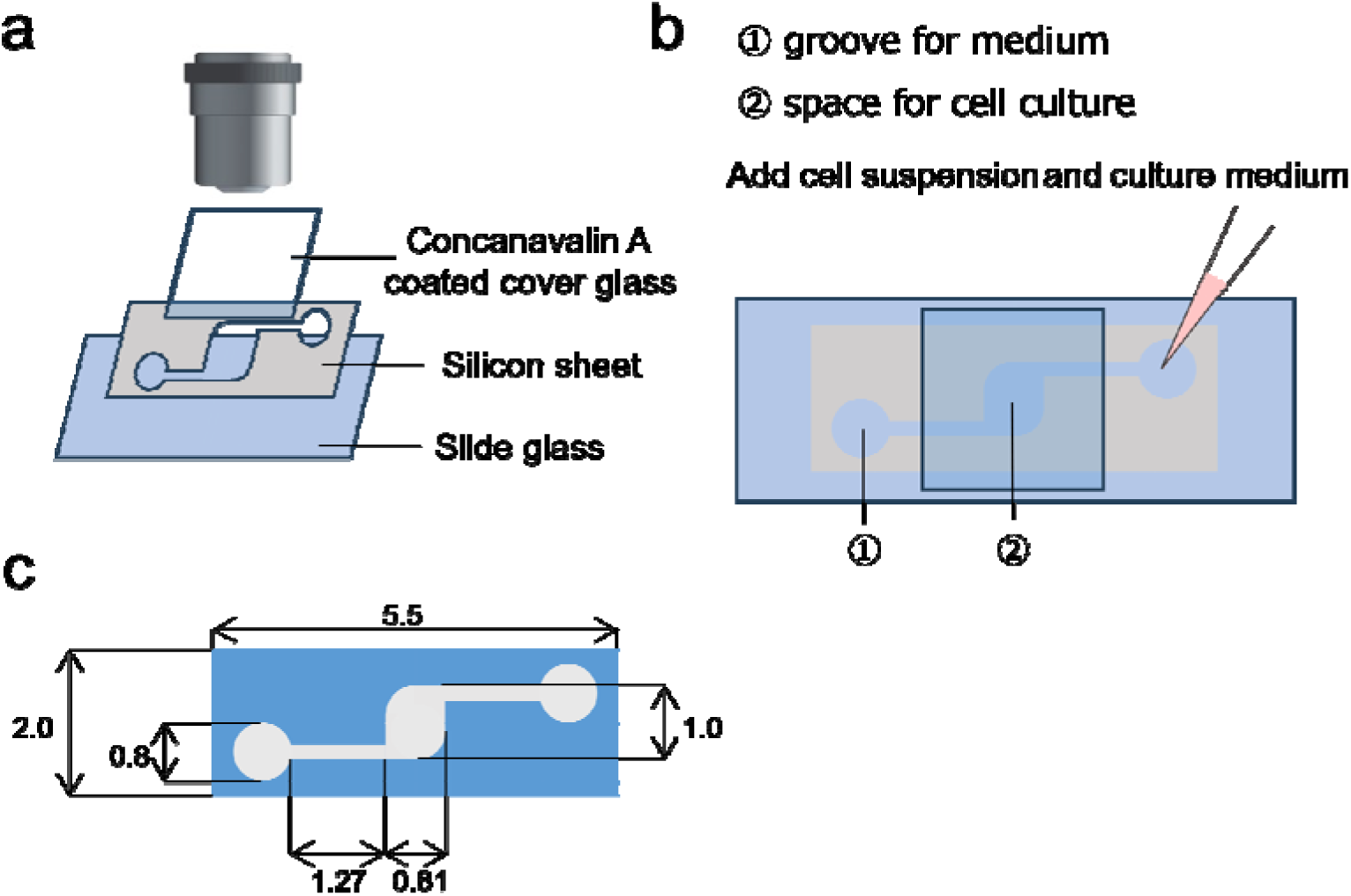
Layout of an observation chamber to obtain time-lapse movies of macrophages with a higher resolution. **a,** Schemes for the assembly of the apparatus and the observation of macrophages. A silicon sheet was attached to a slide grass. A concanavalin-coated cover glass was placed on the top of the silicon sheet. **b,** The suspension of macrophages was added from the groove for the medium, and the culture medium was further supplemented to adapt the desiccation of the chamber. **c,** The blue script of a silicon sheet showing the clipping size (cm).

**Extended Data Fig. 5.**
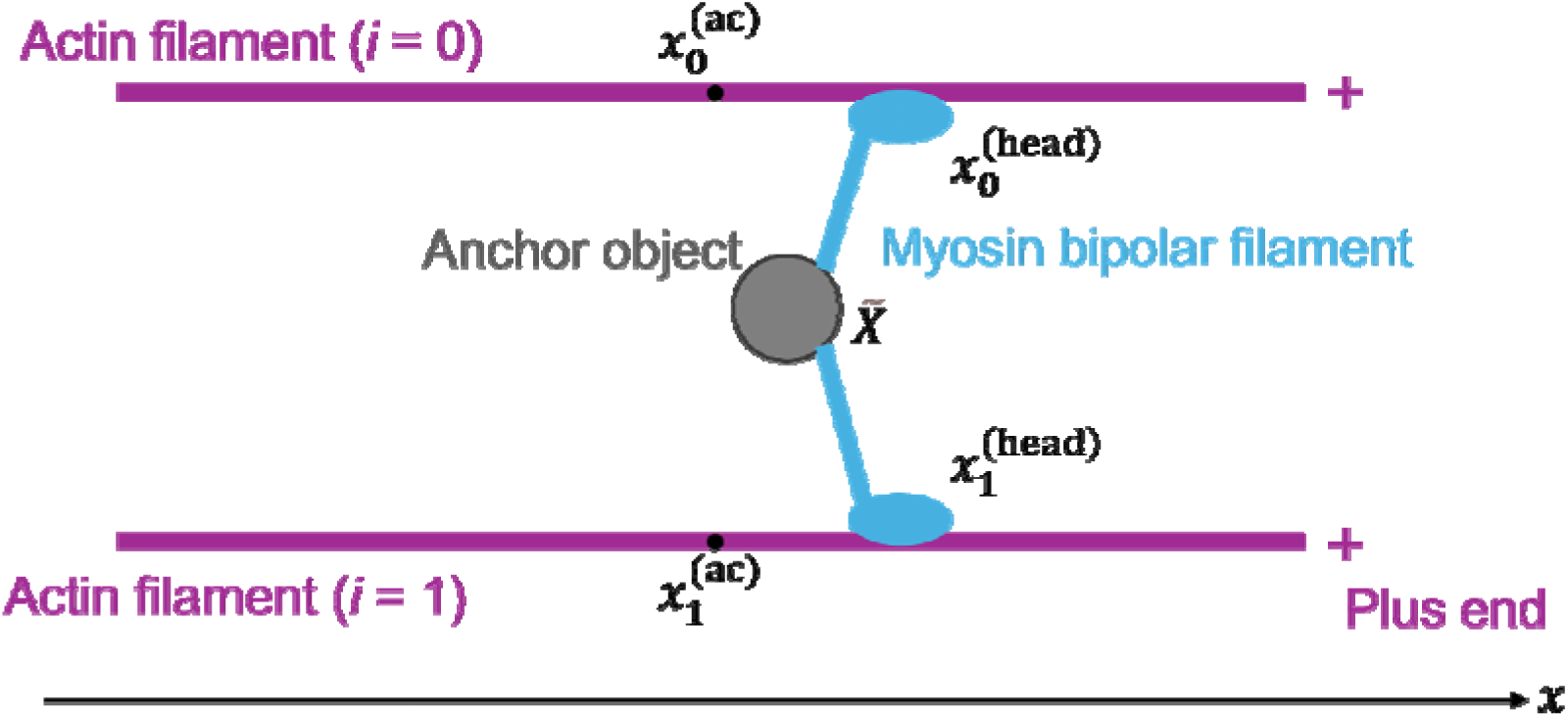
Schematic illustration of the mathematical model. The model considers two parallel actin filaments (indexed by i = 0,1) undergoing translational motion only. A myosin bipolar filament, anchored at its center to an object such as cytoplasmic structure, is positioned between the actin filaments. Each end of the myosin filament has a head domain representing multiple myosin heads, which move toward the plus-end (indicated by + symbols) of the respective actin filament at a constant velocity v. Detailed equations of motion and simulation parameters are provided in the main text and Table 7.

**Extended Data Fig. 6.**
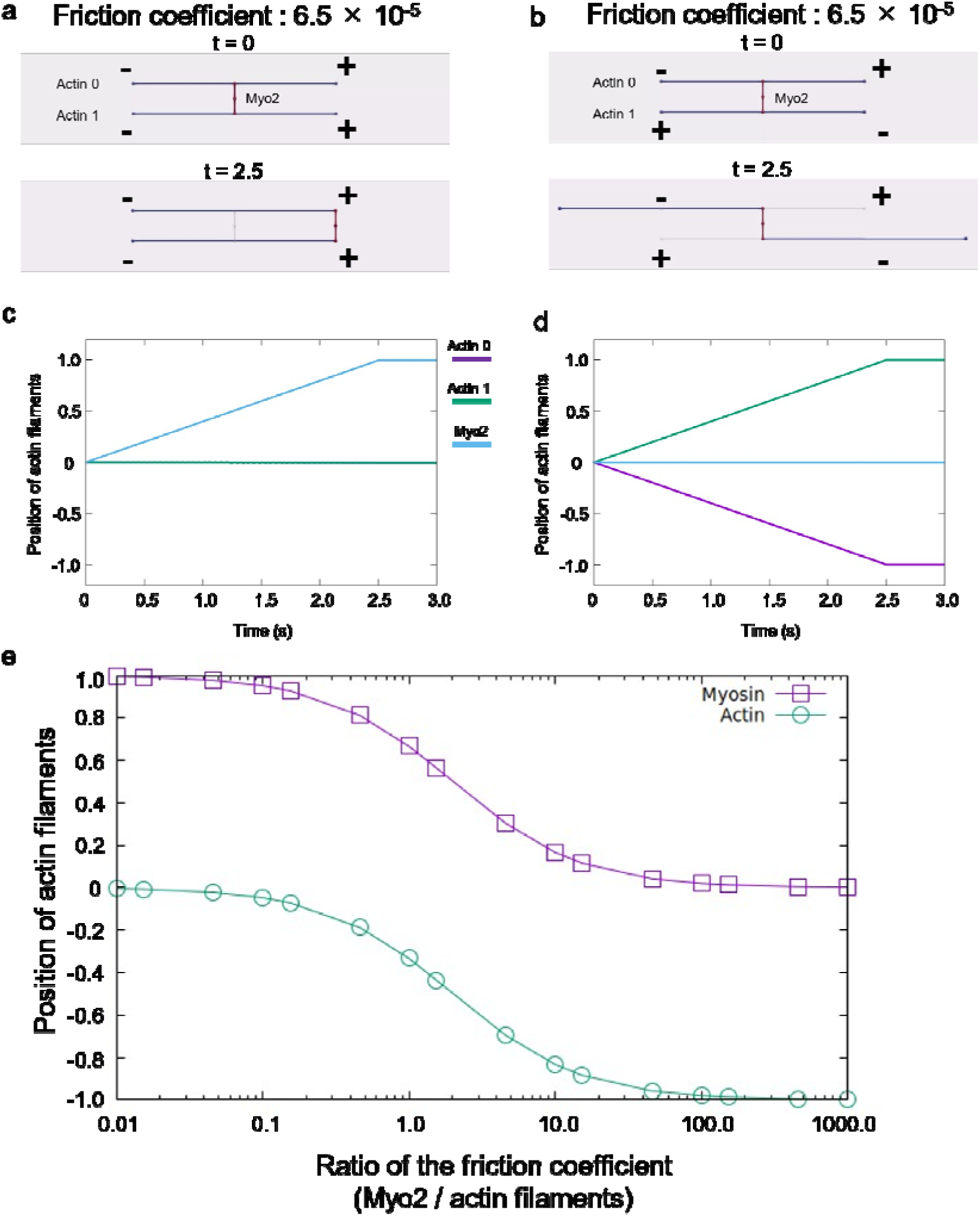
Mathematical simulations to exhibit the translocation of F-actin by Myo2. **a** and **b**, Simulations of translocation of two actin filaments under the condition of friction coefficient for Myo2 (6.5 × 10^−5^). Actin filaments are parallel (**a**) and antiparallel (**b**). t indicates the time point. Symbol – and + mean minus (pointed) and plus (barbed) end of actin filaments, respectively. **c** and **d**, The graphs represent the positional changes of actin filaments over time. **c** and **d** are the result of simulation (**a**) and (**b**), respectively. Blue, green, and magenta lines indicate the positional changes of Myo2 and each of the two actin filaments, respectively. Magenta line (Actin 0) and green line (Actin 1) overlap (c). **e**, Relationship between the displacement at the end of the simulation and the ratio of the friction coefficient of Myo2 and actin filaments. Green and magenta lines indicate the results of actin filaments and Myo2, respectively.

**Supplementary Video 1**

A time-lapse video of a wild type macrophage expressing *Lifeact-mCherry* for 6 h: Lifeact-mCherry is colored magenta. Scale bar is 10 μm.

**Supplementary Video 2**

Time-lapse videos of macrophages misexpressing *Lifeact-mGFP6*; wild type, *Myo1D* mutant, *Myo1D* mutant overexpressing *Myo1D* OE, wild type overexpressing *Myo1D*, wild type overexpressing *Myo1C* for 10 min. Lifeact-mGFP6 is colored green. Scale bars are 5 μm.

**Supplementary Video 3**

A time-laps video of a wild type macrophage misexpressing *Cnn-GFP* and *Redstinger* for 6 h. Cnn-GFP and Redstinger are colored green and magenta, respectively. Scale bar is 10 μm.

**Supplementary Video 4**

Time-laps videos of fluorescent-labeled F-actin in the modified *in vitro* motility assay involving Myo1D (left panel) and Myo1C (right panel). The videos are displayed at 30 × speed.

**Supplementary Video 5**

Simulation video showing the movement of actin filaments using a mathematical model controlled by the friction coefficient of Myo2.

